# Reconciling S-LDSC and LDAK models and functional enrichment estimates

**DOI:** 10.1101/256412

**Authors:** Steven Gazal, Carla Marquez-Luna, Hilary K. Finucane, Alkes L. Price

## Abstract

Recent work has highlighted the importance of accounting for linkage disequilibrium (LD)-dependent genetic architectures in analyses of heritability, motivating the development of the baseline-LD model used by stratified LD score regression (S-LDSC) and the LDAK model. Although both models include LD-dependent effects, they produce very different estimates of functional enrichment (with larger estimates using the baseline-LD model), leading to different interpretations of the functional architecture of complex traits. Here, we perform formal model comparisons and empirical analyses to reconcile these findings. First, by performing model comparisons using a likelihood approach, we determined that the baseline-LD model attains likelihoods across 16 UK Biobank traits that are substantially higher than the LDAK model. Second, we determined that S-LDSC using a combined model (unlike methods that use the LDAK or baseline-LD models) produces robust enrichment estimates in simulations under both the LDAK and baseline-LD models, validating the combined model as a gold standard. Third, in analyses of 16 UK Biobank traits, we determined that enrichment estimates obtained by S-LDSC using the combined model were nearly identical to those obtained by S-LDSC using the baseline-LD model (concordance correlation coefficient *ρ_c_* = 0.99), but were larger than those obtained using LDAK (*ρ_c_* = 0.54). Notably, LDAK enrichment estimates were much higher for a non-default version of LDAK that models SNPs in perfect LD differently by assigning non-zero weights to all SNPs. Our results support the use of the baseline-LD model and confirm the existence of functional annotations that are highly enriched for complex trait heritability.

## Introduction

Recent work has highlighted the importance of accounting for linkage disequilibrium (LD)-dependent genetic architectures in analyses of heritability^1-4^. Two models incorporating LD-dependent architectures have been proposed for analyses of functional enrichment: the baseline-LD model^4^ used by stratified LD score regression^4,5^ (S-LDSC), and the LDAK model^1,3^. Although both models include LD-dependent effects, they produce very different estimates of functional enrichment (e.g. 9.35x (±0.80) in ref. ^4^ and 1.34x (±0.26) in ref. ^3^ for conserved regions), leading to different interpretations of the functional architecture of complex traits.

Here, we perform three sets of analyses, incorporating a combined model, to reconcile these findings. First, we perform formal model comparisons across 16 UK Biobank traits^6,7^ using a likelihood approach^3^. Second, we compare methods for estimating functional enrichment using these models in simulations under each model. Third, we apply these methods for estimating functional enrichment to 16 UK Biobank traits. We find that the baseline-LD model and the combined model attain the highest likelihoods in all model comparisons, and that the combined model produces robust results in simulations under each model and results similar to the baseline-LD model across 16 UK Biobank traits. We also find that lower enrichment estimates for LDAK could potentially be explained by the assignment of zero weights to most SNPs.

## Results

### The GCTA, LDAK and baseline-LD models

We define several heritability models, including the “GCTA”, LDAK and baseline-LD models. We define each model according to *Var*(*β_j_*), the variance of normalized causal effect sizes *β_j_* for each SNP *j*. We note that *Var*(*β_j_*) is equal to the variance of per-allele causal effect sizes times 2*p_j_*(1 – *p_j_*) where *p_j_* is minor allele frequency (MAF), and can also be interpreted as the expected causal per-SNP heritability of SNP *j*. The eight main heritability models defined by Equations (1)-(8) below are summarized in Table 1.

We first consider an infinitesimal model^8^ in which normalized causal effect size variances are constant:

**Table 1:**
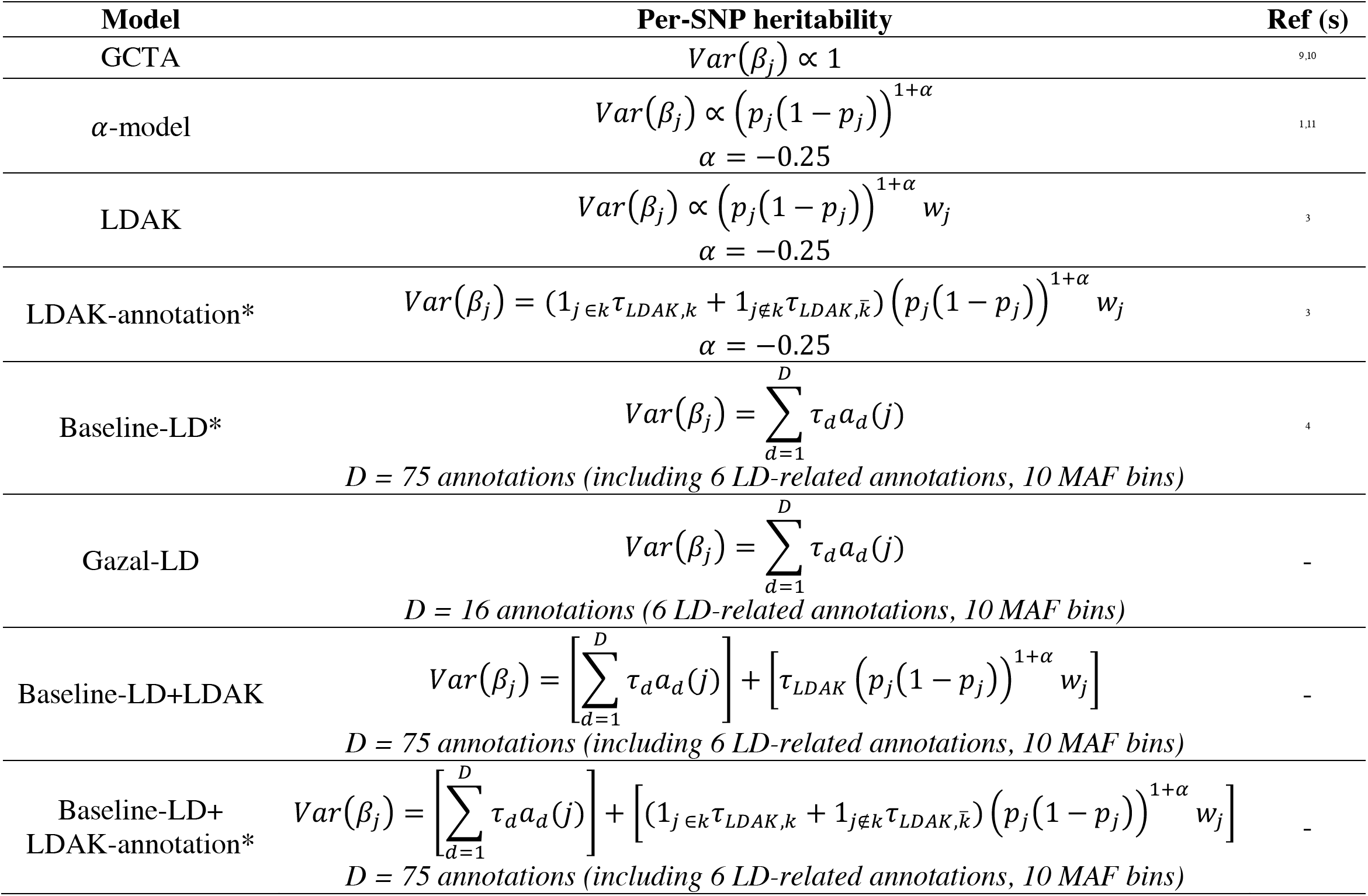
Models of per-SNP heritability. We summarize the eight main heritability models defined by Equations (1)-(8). We define each model according to *Var*(*β_j_*), the variance of normalized causal effect sizes *β_j_* for each SNP *j*. * denotes models that are used to estimate functional enrichment in this manuscript.

| Model | Per-SNP heritability | Ref (s) |
| --- | --- | --- |
| GCTA | $Var(\beta_j) \propto 1$ | 9,10 |
| α-model | $Var(\beta_j) \propto (p_j(1-p_j))^{1+\alpha}$<br>$\alpha = -0.25$ | 1,11 |
| LDAK | $Var(\beta_j) \propto (p_j(1-p_j))^{1+\alpha} w_j$<br>$\alpha = -0.25$ | 3 |
| LDAK-annotation* | $Var(\beta_j) = (1_{j \in k} \tau_{LDAK,k} + 1_{j \notin k} \tau_{LDAK,\bar{k}}) (p_j(1-p_j))^{1+\alpha} w_j$<br>$\alpha = -0.25$ | 3 |
| Baseline-LD* | $Var(\beta_j) = \sum_{d=1}^D \tau_d a_d(j)$<br>*D = 75 annotations (including 6 LD-related annotations, 10 MAF bins)* | 4 |
| Gazal-LD | $Var(\beta_j) = \sum_{d=1}^D \tau_d a_d(j)$<br>*D = 16 annotations (6 LD-related annotations, 10 MAF bins)* | - |
| Baseline-LD+LDAK | $Var(\beta_j) = \left[ \sum_{d=1}^D \tau_d a_d(j) \right] + \left[ \tau_{LDAK} (p_j(1-p_j))^{1+\alpha} w_j \right]$<br>*D = 75 annotations (including 6 LD-related annotations, 10 MAF bins)* | - |
| Baseline-LD+LDAK-annotation* | $Var(\beta_j) = \left[ \sum_{d=1}^D \tau_d a_d(j) \right] + \left[ (1_{j \in k} \tau_{LDAK,k} + 1_{j \notin k} \tau_{LDAK,\bar{k}}) (p_j(1-p_j))^{1+\alpha} w_j \right]$<br>*D = 75 annotations (including 6 LD-related annotations, 10 MAF bins)* | - |

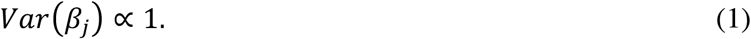

This model was called the “GCTA model” in ref. ^3^ (as it was implemented in the original version of the GCTA software^9,10^), and we adhere to that convention here; we note that models incorporating LD bins (and/or MAF bins) have also been implemented in the GCTA software^2,11^, but we do not consider those models here as they have not been used for estimating functional enrichment. We note that the S-LDSC method does not use or assume the GCTA model (see Methods for estimating functional enrichment).

We next consider a model in which MAF-dependent architectures are modeled using an *α* parameter^1,11^:

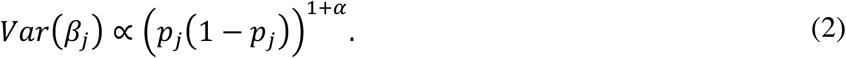

We refer to this model as the “*α*-model”. We note that the α-model with *α* = −1.00 is identical to the GCTA model, and that recent work has estimated mean *α* values of −0.38 to −0.36 for UK Biobank traits^12,13^.

Speed et al. have made an important contribution to the literature by highlighting the potential ramifications of LD-dependent architectures^1,3^. They recently introduced an “LDAK model” that models both LD-dependent and MAF-dependent architectures^3^:

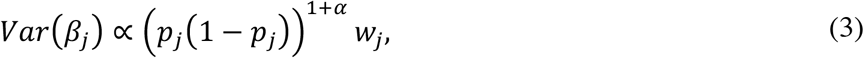

where *w_j_* denote LDAK weights reflecting smaller expected causal per-SNP heritability for high-LD SNPs; for example, *“two SNPs in perfect LD are expected to contribute only half the heritability of two SNPs showing no LD*” (p.1 of ref. ^3^; also see Figure S1). The weights *w_j_* are fixed based on LD patterns, and are not estimated using trait data. We fix *α* = −0.25 in all analyses using the LDAK model, as recommended^3^. (We do not consider an earlier version of the LDAK model^1^ with *α* = −1.00.) For consistency and ease of comparison, we also fix *α* = −0.25 in all analyses using the *α*-model.

Ref. ^3^ also proposed a model, which we call the “LDAK-annotation model”, that extends the LDAK model to incorporate a single functional annotation *k*:

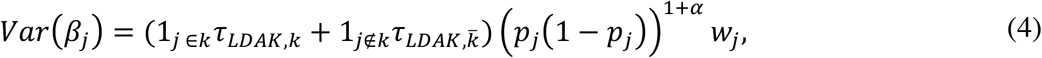

where 1_*j*∈*k*_ and 1_*j*∉*k*_ are indicator functions, and 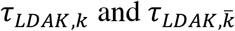 are coefficients estimated using trait data.

We recently introduced a “baseline-LD” model that models LD-dependent, MAF-dependent and functional architectures^4^:

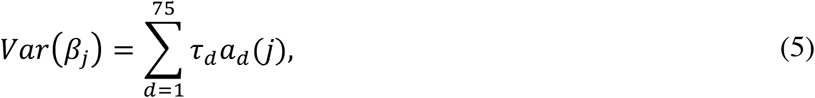

where the coefficients *τ_d_* denote conditional contributions of annotations *a_d_* to expected per-SNP heritability. The coefficients *τ_d_* are estimated using trait data. The 75 annotations *a_d_* include 6 continuous-valued LD-related annotations, 10 common SNP MAF bins, and overlapping functional annotations (e.g. coding, conserved, regulatory) from a previous “baseline model”^5^ (see Methods and Table S1). We note that the baseline-LD model jointly models many overlapping annotations to provide flexibility and robustness to model misspecification^5^.

For comparison purposes, we define a “Gazal-LD model” in which Equation (5) is restricted to the 6 LD-related annotations and 10 common SNP MAF bins from the baseline-LD model:

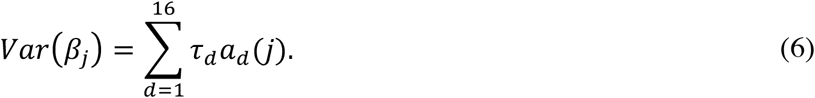

We also define a “baseline-LD+LDAK model” that combines the baseline-LD (Equation (5)) and LDAK (Equation (3)) models:

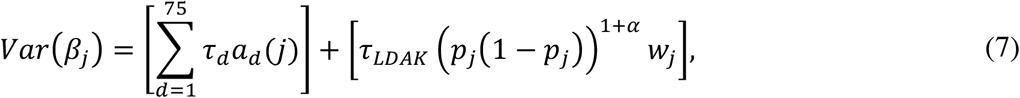

where coefficients *τ_d_* and *τ_LDAK_* are estimated using trait data. This model provides maximum flexibility to fit the data effectively, e.g. either if the data follows the baseline-LD model or if the data follows the LDAK model, through the estimation of the coefficients *τ_d_* and *τ_LDAK_* from trait data.

Finally, we define a “baseline-LD+LDAK-annotation model” that combines the baseline-LD (Equation (5)) and LDAK-annotation models for a single functional annotation *k* (Equation (4)) present in the baseline-LD model:

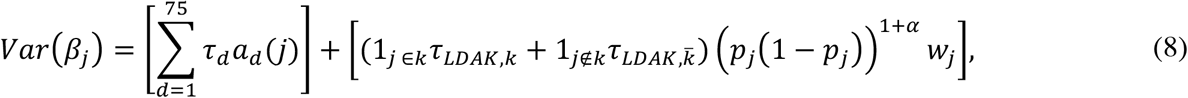

where coefficients *τ_d_*, *τ_LDAK,k_* and 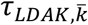 are estimated using trait data.

We note several differences between the LDAK and baseline-LD models. First, the LDAK model has zero degrees of freedom parameterizing the relationship between LD and *Var*(*β_j_*), whereas the baseline-LD model has six degrees of freedom. Second, the LDAK model has one degree of freedom parameterizing the relationship between MAF and *Var*(*β_j_*) (or zero degrees of freedom when fixing *α* = −0.25 as recommended), whereas the baseline-LD model has 10 degrees of freedom. Third, the LDAK model assumes that LD and MAF effects combine multiplicatively, whereas the baseline-LD model assumes that they combine additively. Fourth, the LDAK model assumes the same architecture for every trait (when *α* is fixed to the recommended value), whereas the baseline-LD model allows for trait-specific architectures. Fifth, the baseline-LD model includes functional annotations, unlike the LDAK model (and Gazal-LD model). Finally, we note that because these two models have different numbers of degrees of freedom (75 vs. 1), the baseline-LD model requires larger data sets to estimate its parameters (see below), whereas the LDAK model can be applied to smaller data sets.

### Comparison of GCTA, LDAK and baseline-LD models using likelihood approach

We compared the 6 models defined by Equations (1)-(3) and (5)-(7) using the likelihood approach of ref. ^3^. (We excluded the LDAK-annotation model defined by Equation (4) and the baseline-LD+LDAK-annotation model defined by Equation (8), as these models focus on a single functional annotation.) We analyzed *N* = 20K unrelated British-ancestry samples, due to LDAK computational constraints. When computing likelihoods for the baseline-LD, Gazal-LD and baseline-LD+LDAK models, we estimated all *τ* parameters out-of-sample by running S-LDSC on the remaining unrelated British-ancestry UK Biobank samples, excluding the *N* = 20K samples and samples related to those samples (average *N* = 299K); when computing likelihoods for the GCTA model, α-model and LDAK model, we did not estimate any parameters out-of-sample. All six models include one heritability parameter that is maximized in-sample when estimating the likelihood. We note that these formal model comparisons evaluate the respective *models*, irrespective of any *method* for estimating functional enrichment. The authors of ref. ^3^ reported that the LDAK model (with *α* = −0.25) attained a higher likelihood than both the GCTA model and the *α*-model (with *α* = −0.25), on average across 42 traits. We computed likelihoods for 16 independent UK Biobank traits^6,7^ (see Methods and Table S2). We analyzed either genotypes imputed using the 1000 Genomes (1000G) reference panel^14^ (as in ref. ^3^; *M* = 2.8 million well-imputed SNPs) or genotypes imputed using the Haplotype Reference Consortium (HRC) reference panel^15^ (*M* = 4.6 million well-imputed SNPs); in each case, we restricted our analyses to SNPs with INFO score^16^ ≥ 0.99 and MAF ≥ 1% (as in ref. ^3^), and excluded variants in the MHC region. We note that well-imputed HRC SNPs (median LD score = 110) have lower LD than well-imputed 1000G SNPs (median LD score = 139; Figure S2) due to larger reference panel sample size. The number of SNPs with non-zero LDAK weights was 308K for the 1000G SNPs and 708K for the HRC SNPs; the remaining 85-89% of SNPs have *w_j_* = 0, so that *Var*(*β_j_*) = 0 in Equation (3). (For example, given multiple SNPs in perfect LD, the default version of LDAK will randomly assign a weight of 1 to one of the SNPs and 0 to the others, independent of their functional annotations.) The assignment of zero weights to most SNPs may have little impact on likelihood computations (in which a SNP in perfect LD with a zero-weight SNP can act as a proxy), but may have a substantial impact on functional enrichment, in which an out-of-annotation SNP in perfect LD with a zero-weight in-annotation SNP cannot act as a proxy (see Impact of proportion of SNPs with zero weights on functional enrichment estimates).

We first analyzed *M* = 2.8 million well-imputed 1000G SNPs (Figure 1a and Table S3). We report the change in log likelihood compared to the LDAK model (ΔLL) for each other model compared, summed across the 16 traits. We confirmed that the GCTA model and the *α*-model attained lower likelihoods than the LDAK model (ΔLL ≤ −220; ΔLL < 0 for most traits), consistent with ref. ^3^. On the other hand, the Gazal-LD, baseline-LD and baseline-LD+LDAK models attained substantially higher likelihoods than the LDAK model (ΔLL ≥ 439; ΔLL > 0 for most traits).

**Figure 1:**
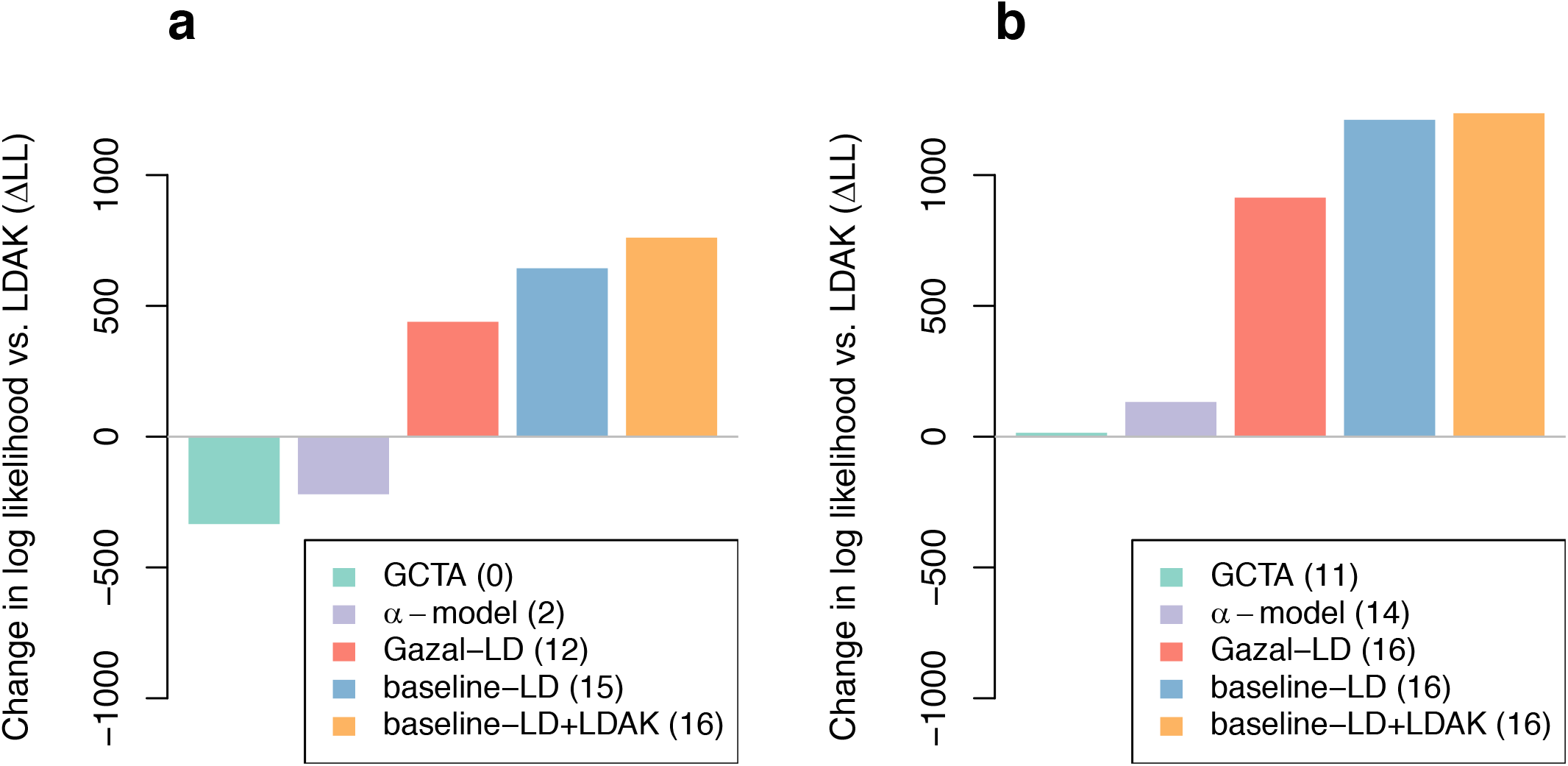
Likelihood comparison of different models of per-SNP heritability. We report the change in log likelihood compared to the LDAK model (ΔLL) of five other per-SNP heritability models, summed across 16 independent UK Biobank traits (*N* = 20K). All six models include one heritability parameter that is maximized in-sample when estimating the likelihood. (**a**) Analyses using *M* = 2,835,699 well-imputed 1000G SNPs (as in ref. ^3^). (**b**) Analyses using *M* = 4,631,901 well-imputed HRC SNPs. Numbers between parentheses in figure legends indicate the number of traits with ΔLL > 0. Numerical results, including results for each trait, are reported in Table S3. The *M* = 4.6 million well-imputed HRC SNPs consistently attained higher likelihoods than the *M* = 2.8 million well-imputed 1000G SNPs in comparisons using the same model (Table S3).

We then analyzed *M* = 4.6 million well-imputed HRC SNPs (Figure 1b and Table S3). In this analysis, the GCTA model and the *α*-model attained slightly higher likelihoods than the LDAK model (ΔLL ≥ 15-133; ΔLL > 0 for most traits). In addition, the Gazal-LD, baseline-LD and baseline-LD+LDAK models attained much higher likelihoods than the LDAK model (ΔLL ≥ 914; ΔLL > 0 for 16/16 traits in each case). The baseline-LD and baseline-LD+LDAK models attained similar likelihoods (ΔLL = 1211 and ΔLL = 1236, respectively), implying that LDAK weights provide limited additional information conditional on baseline-LD annotations. We obtained similar results in analyses using the baseline model of ref. ^5^, in analyses with sample size *N* = 7K, analogous to ref. ^3^, as well as in analyses where we trained model parameters using a meta-analysis across 16 traits (Figure S3). We also obtained similar conclusions using out-of-sample polygenic prediction^17^, as proposed in ref. ^18^ (see Methods and Figure S4; 6.2 million UK10K SNPs, as in ref. ^17^).

In summary, we reached two main conclusions. First, the baseline-LD model and the combined model consistently attained much higher likelihoods than the GCTA model, *α*-model or LDAK model. Second, the GCTA model, *α*-model and LDAK model attained relatively similar likelihoods, although the likelihood order depended on the SNP set. We focused all subsequent analyses on the *M* = 4.6 million well-imputed HRC SNPs, which attained higher likelihoods than *M* = 2.8 million well-imputed 1000G SNPs in comparisons using the same model (ΔLL = 860-1427; Table S3).

### Methods for estimating functional enrichment

We use three models to estimate functional enrichment: the baseline-LD model (Equation (5)), the LDAK-annotation model (Equation (4)), and the baseline-LD+LDAK-annotation model (Equation (8)). We only consider models with LD-dependent architectures, as it is widely recognized that failure to model LD-dependent architectures can lead to biased enrichment estimates^3,4^. We define four main methods for estimating functional enrichment using these three models. Each functional enrichment method is defined by its model, inference procedure, and enrichment estimand; thus, there does not exist a 1-1 correspondence between models and functional enrichment methods. The four main methods are summarized in Table 2. Further details are provided in the Methods section.

**Table 2:** Main methods for estimating functional enrichment. We summarize the four main methods for estimating functional enrichment. For each method, we list their input data, per-SNP heritability model, inference method, estimand and SNP set (either the set of reference SNPs used to compute LD scores for methods using the S-LDSC inference method, or the set of SNPs included in the GRM for methods using the REML inference method). Methods that analyze summary statistics (SS) were applied to summary statistics computed using *N* = 434K individuals; methods that analyze raw genotype/phenotype data (Raw) were applied to *N* = 20K individuals (see main text). Seven auxiliary methods are described in Table S5.

| Method | Data | Model | Inference | Estimand | SNP set | Ref |
| --- | --- | --- | --- | --- | --- | --- |
| S-LDSC | SS | Baseline-LD | S-LDSC | S-LDSC | $AC \geq 5$<br>($M = 13.3\text{M UK10K SNPs}$) | <sup>4</sup> |
| LDAK | Raw | LDAK-annotation | REML | LDAK | $MAF \geq 1\%$, $INFO \geq 0.99$<br>($M = 4.6\text{M HRC SNPs}$) | <sup>3</sup> |
| S-LDSC+LDAK<br>(S-LDSC estimand) | SS | Baseline-LD+<br>LDAK-annotation | S-LDSC | S-LDSC | $AC \geq 5$<br>($M = 13.3\text{M UK10K SNPs}$) | - |
| S-LDSC+LDAK<br>(LDAK estimand) | SS | Baseline-LD+<br>LDAK-annotation | S-LDSC | LDAK | $MAF \geq 1\%$<br>($M = 7.7\text{M UK10K SNPs}$) | - |

SS: summary statistics; AC: allele count; INFO: INFO score.

The first method, which we call the “S-LDSC method”, infers the parameters of the baseline-LD model (Equation (5)) with S-LDSC^4,5^ using default settings. S-LDSC infers model parameters by regressing chi-square statistics on partitioned LD scores, and only requires summary statistics and an LD reference panel. In all analyses below, we use UK10K as the LD reference panel. The method uses all reference panel SNPs with minor allele count ≥ 5, as previously recommended^4,5^. The method uses the “S-LDSC estimand”: the enrichment of annotation *d* is defined as the proportion of heritability causally explained by common SNPs (MAF ≥ 5%) in annotation *d* divided by the proportion of common SNPs in the annotation^4,5^. The method should only be applied to data sets with highly significant total heritability (e.g. z-score > 7 for total heritability), as previously recommended^5^; this generally requires a sample size of *N* ≥ 50K samples, as with all summary statistic based methods that we consider in this study. We note that we previously recommended^4^ that functional enrichment analyses using S-LDSC should be performed using the baseline-LD model^4^ instead of our earlier baseline model^5^, which produces slightly inflated estimates (see Supplementary Table 8c of ref. ^4^); we also note that analyses using two-category S-LDSC (i.e. extending the GCTA model to two categories) are not recommended due to model misspecification effects of unmodeled annotations (see Figure 2 of ref. ^5^).

**Figure 2:**
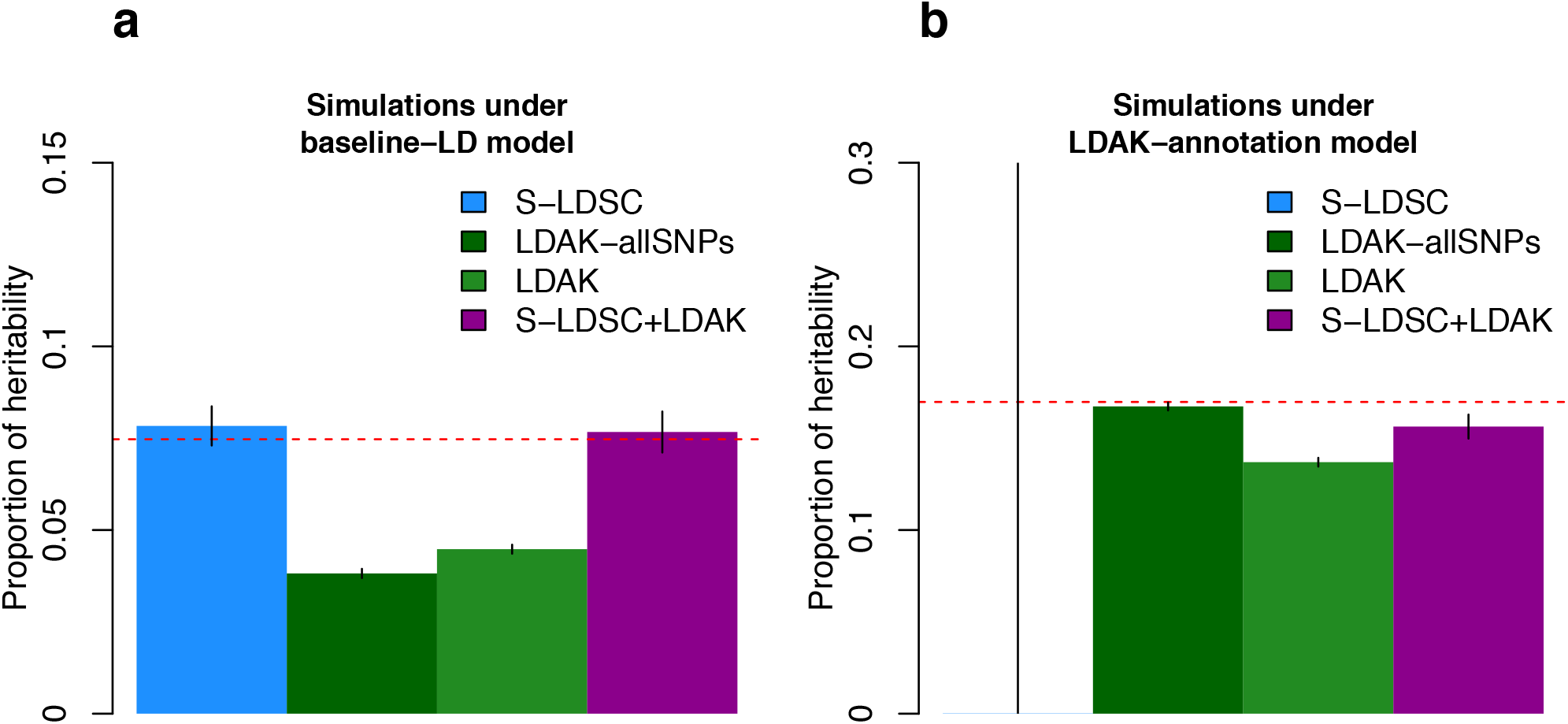
Simulations with coding enrichment under the baseline-LD and LDAK-annotation models. We report the estimated proportion of heritability explained by coding variants in simulations in which we simulated effect sizes based on the baseline-LD model (**a**) or based on coding enrichment under the LDAK-annotation model (**b**). The S-LDSC method uses S-LDSC with the baseline-LD model. The LDAK-allSNPs method analyzes all SNPs with MAF ≥1%. The LDAK method analyzes all SNPs with MAF ≥1% and INFO score ≥ 0.99 (as recommended^3^). The S-LDSC+LDAK method uses S-LDSC with the baseline-LD+LDAK-annotation model. We report the proportion of heritability explained rather than enrichment, due to the different LDAK and S-LDSC enrichment estimands. Dashed red lines indicate the true simulated value. Results are averaged across 500 simulations. Error bars represent 95% confidence intervals. Results for other simulation scenarios and other methods are reported in Figure S5. Numerical results are reported in Table S6.

The second method, which we call the “LDAK method”, infers the parameters of the LDAK-annotation model (Equation (4)) using restricted maximum likelihood^9,10^ (REML), using default settings as previously described^3^. The method requires raw genotypes/phenotypes. Analyses are restricted to well-imputed SNPs (INFO score ≥ 0.99) with MAF ≥ 1%, as recommended for functional enrichment analyses^3^. The method uses the “LDAK estimand”: the enrichment of annotation *d* is defined as the proportion of heritability causally explained by all SNPs in annotation *d* divided by the proportion of heritability *expected under the LDAK model*^3^. We note the difference between the S-LDSC estimand and the LDAK estimand; although either choice of estimand can be justified, direct comparisons of functional enrichment estimates based on different estimands may yield incorrect conclusions for LD-related annotations (see Supplementary Note). The LDAK method can only be applied to relatively small data sets (e.g. *N* = 20K samples), due to computational constraints of LDAK software.

The third method infers the parameters of the baseline-LD+LDAK-annotation model (Equation (8)) using S-LDSC. The method only requires summary statistics and an LD reference panel. The method uses all reference panel SNPs with minor allele count ≥ 5. Parameters and functional enrichment are estimated using S-LDSC with default parameters. The method uses the S-LDSC estimand. We refer to this method as the “S-LDSC+LDAK (S-LDSC estimand) method”.

The fourth method is identical to the third method, except that the method uses the LDAK estimand and the reference panel SNPs is restricted to all SNPs with MAF ≥ 1% (consistent with the LDAK method). We refer to this method as the “S-LDSC+LDAK (LDAK estimand) method”.

In simulations (see below), we focus on estimating the proportion of heritability explained by SNPs with MAF ≥ 1%, instead of functional enrichment. In this case, we restricted all LD reference panels to SNPs with MAF ≥ 1%, such that there is no difference between the S-LDSC and LDAK estimands. The latter two methods are thus collapsed into a single method, the “S-LDSC+LDAK method”.

### Simulations

We compared methods for estimating functional enrichment by performing simulations using real UK Biobank genotypes and phenotypes simulated under either the baseline-LD model or the LDAK-annotation model. To facilitate comparisons between methods, we restricted the UK Biobank data (and UK10K reference panel data) to SNPs with MAF ≥ 1%, and estimated the proportion of heritability explained by MAF ≥ 1% SNPs in an annotation (instead of estimating functional enrichment using the S-LDSC estimand or LDAK estimand (see Table S4 for a description of the set of SNPs used by S-LDSC, LDAK and S-LDSC+LDAK in our simulations and analyses of UK Biobank traits)). We restricted our simulations to 10,000 unrelated samples, due to the computational limitations of LDAK software. We sampled integer-valued genotypes from UK Biobank imputed dosages, restricting to 578,876 SNPs on chromosome 1 with MAF ≥ 1%. We set trait heritability to 0.5; concentrating this amount of heritability onto a single chromosome enabled highly significant total heritability (z-score > 7) with only 10,000 samples.

We simulated causal effect sizes using either (a) the baseline-LD model with previously estimated parameters^4^, including coding enrichment or (b) coding enrichment under the LDAK-annotation model (see Methods). We compared the S-LDSC, LDAK and S-LDSC+LDAK methods (as noted above, there is no difference between S-LDSC+LDAK (S-LDSC estimand) and S-LDSC+LDAK (LDAK estimand) in these simulations). We also considered an auxiliary method that is identical to the LDAK method except that analyses include all SNPs with MAF ≥ 1% (LDAK-allSNPs), instead of restricting to well-imputed SNPs with MAF ≥ 1% (Table S5). (We do not consider LDAK-allSNPs in our analyses of real UK Biobank traits, because ref. ^3^ recommends that functional enrichment analyses using LDAK should analyze well-imputed SNPs only and because it is unclear how dosage data from poorly imputed SNPs would be incorporated; this complexity does not apply to our simulations, which use integer-valued genotypes both for simulating phenotypes and for inference.)

Results are reported in Figure 2 and Table S6. As expected, S-LDSC and LDAK-allSNPs were unbiased when we simulated coding enrichment under their corresponding models. S-LDSC produced unstable enrichment estimates under the LDAK model, although S-LDSC with constrained intercept produced stable (conservative) enrichment estimates (Figure S5; see Supplementary Note). This inconsistency between S-LDSC and S-LDSC with constrained intercept was not observed in analyses of real traits (see below). (We note that S-LDSC with constrained intercept with the baseline model^5^ instead of the baseline-LD model produced upward biased instead of conservative enrichment estimates (Figure S5), underscoring the importance of modeling LD-dependent architectures.) The LDAK method was downward biased under the LDAK model, as it restricted analyses to well-imputed SNPs, and even more downward biased under the baseline-LD model. In null simulations under the LDAK model (i.e. no functional enrichment), the LDAK method was upward biased, unlike LDAK-allSNPs and S-LDSC+LDAK (Figure S5). Notably, S-LDSC+LDAK was unbiased in simulations under the baseline-LD model, with an extremely small downward (conservative) bias in simulations under the LDAK model. We thus defined the S-LDSC+LDAK method as a “gold standard” for estimating the proportion of heritability, allowing us to define the S-LDSC+LDAK (S-LDSC estimand) and S-LDSC+LDAK (LDAK estimand) methods as gold standards for estimating the S-LDSC and LDAK functional enrichment estimands, respectively.

We obtained similar conclusions in simulations with different numbers of causal variants, showing that the methods are robust to less sparse architectures (Figure S6). We also obtained similar conclusions when using conserved and DNase I hypersensitive sites (DHS) enrichment (Figure S5); however, we note that LDAK was even more strongly downward biased for conserved enrichment simulated under the baseline-LD model. Finally, we also determined that S-LDSC+LDAK was unbiased in simulations under the baseline-LD+LDAK-annotation model (see Methods and Figure S5).

### Functional enrichment estimates across 16 UK Biobank traits

We compared the S-LDSC and LDAK methods to the “gold standard” S-LDSC+LDAK (S-LDSC estimand) and S-LDSC+LDAK (LDAK estimand) methods, respectively, by meta-analyzing their enrichment estimates for 28 main functional annotations (Table S1) across the 16 UK Biobank traits^6,7^ (see Methods and Table S2). We applied S-LDSC, S-LDSC+LDAK (S-LDSC estimand) and S-LDSC+LDAK (LDAK estimand) to summary statistics computed using BOLT-LMM v2.3 (ref. ^19^; see URLs) from HRC-imputed dosage data (average *N* = 434K). We applied LDAK to *N* = 20K samples, analyzing well-imputed HRC SNPs (see above). We used concordance correlation coefficient^20^ (*ρ_c_*) as our comparison metric (see Methods).

S-LDSC and S-LDSC+LDAK (S-LDSC estimand) produced nearly identical estimates of enrichment (*ρ_c_* = 0.99), with a consistent maximum enrichment for conserved regions (8.11x, s.e. = 0.54x and 7.48x, s.e. = 0.49x, respectively; Figure 3a and Table S7). These results imply that adding annotations constructed from LDAK model weights did not significantly change S-LDSC enrichment estimates. We note that enrichment estimates from S-LDSC were nearly identical to enrichment estimates from S-LDSC with constrained intercept (*ρ_c_* = 1.00; Figure S7), unlike in simulations under the LDAK model.

**Figure 3:**
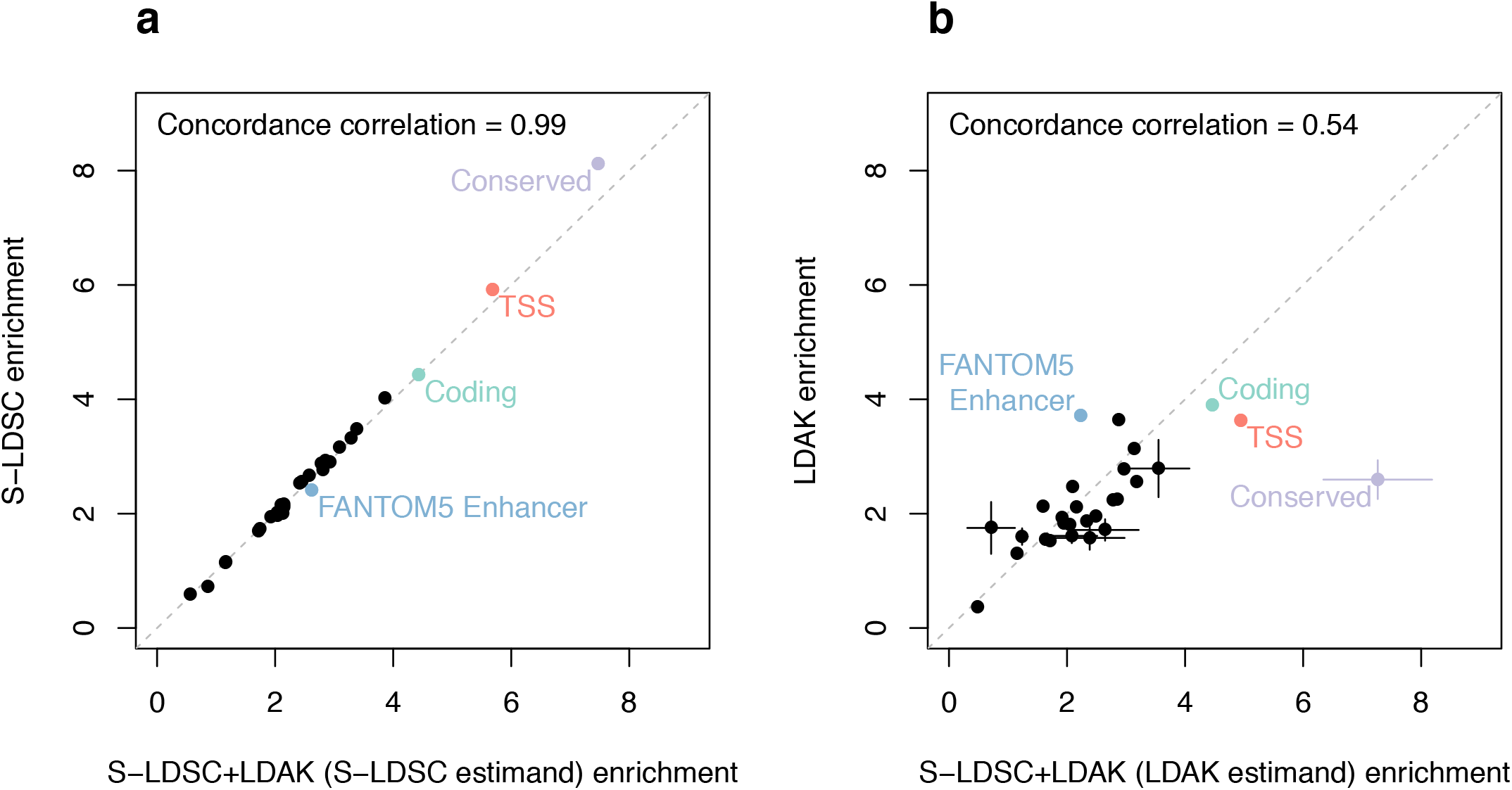
Comparison of functional enrichment estimates in analyses of UK Biobank traits. We report the enrichment of S-LDSC+LDAK (S-LDSC estimand) vs. S-LDSC (using the baseline-LD model) (**a**) and S-LDSC+LDAK (LDAK estimand) vs. LDAK (**b**) for 28 functional annotations, meta-analyzed across 16 independent UK Biobank traits. In each case we report the concordance correlation coefficient *ρ_c_*. Dashed grey lines represent *y* = *x*. Error bars represent 95% confidence intervals for annotations for which the estimated enrichment is significantly different (*P* < 0.05) between the two methods. Numerical results are reported in Table S7.

Enrichment estimates from LDAK were lower than enrichment estimates from S-LDSC+LDAK (LDAK estimand) (*ρ_c_* = 0.54), with 8 annotations having a nominally significant difference in enrichment (*P* < 0.05), including a large difference for conserved regions (2.60x, s.e. = 0.17x and 7.26x, s.e. = 0.47x, respectively; Figure 3b and Table S7); these results are consistent with the LDAK downward bias observed in simulations under the baseline-LD model (Figure 2a). However, we note that the maximum LDAK enrichment (coding regions; 3.90x, s.e. = 0.36x), was larger than the maximum significant LDAK enrichment reported in ref. ^3^ (2.51x, s.e. = 0.34x; Figure S8). We hypothesize that the difference comes from the set of traits investigated, as we observed that our LDAK enrichment estimates were little changed when analyzing a similar sample size and SNPs set as in ref. ^3^ (*N* = 7K individuals and *M* = 2.8M 1000G SNPs; *ρ_c_* = 0.87; Figure S9). Finally, we note that S-LDSC+LDAK (S-LDSC estimand) and S-LDSC+LDAK (LDAK estimand) enrichment estimates were nearly identical (*ρ_c_* = 0.99, no annotation significantly different; Figure S10), showing that the choice of enrichment estimand has little impact on results for our main 28 annotations, although we caution that consistent estimands are essential when comparing results for LD-related annotations (see Supplementary Note). We hypothesized that the large difference between LDAK and S-LDSC+LDAK (LDAK estimand) enrichment estimates for conserved regions could be due to misspecification of the LDAK model. Indeed, we determined that the difference between inference methods (S-LDSC vs. REML), sample sizes (average *N* = 434K vs. *N* = 20K), and/or sets of SNPs (7.7M SNPs with MAF ≥ 1% vs. 4.6M SNPs with MAF ≥ 1% and INFO score ≥ 0.99) did not impact our conclusions (see Supplementary Note).

In summary, we reached three main conclusions. First, S-LDSC enrichment estimates (but not LDAK enrichment estimates) are concordant with the “gold standard” S-LDSC+LDAK method (with the appropriate estimand). Second, LDAK enrichment estimates were higher than in ref. ^3^, with estimates closer to S-LDSC except for conserved regions (Figure S11). Third, lower enrichment estimates for LDAK for conserved regions are likely due to model misspecification, consistent with the lower LDAK likelihood in formal model comparisons (Figure 1) and strong downward bias in LDAK enrichment estimates in simulations under the baseline-LD model (Figure S5; also see Figure 2).

### Impact of proportion of SNPs with zero weights on functional enrichment estimates

It is possible that the LDAK model’s assignment of zero weights (and thus zero heritability) to 85-89% of SNPs (see Comparison of GCTA, LDAK and baseline-LD models using likelihood approach) could reflect a misspecification of the LDAK model, if those SNPs do not actually have zero heritability. If so, this could have a substantial impact on functional enrichment, in which an out-of-annotation SNP in perfect LD with a zero-weight in-annotation SNP cannot act as a proxy. For this reason, we sought to investigate the impact of the proportion of SNPs with zero weights on functional enrichment estimates.

First, we modified S-LDSC with the baseline-LD model by restricting the LD reference panel to the 708K HRC SNPs with non-zero weight in the LDAK model, effectively assigning zero heritability to the remaining SNPs (S-LDSC-nonzeroLDAKSNPs; see Table S5). We determined that S-LDSC enrichment estimates decreased considerably and were generally similar to or lower than LDAK estimates (Figure S12; e.g. similar enrichment estimate for conserved regions: 2.93x, s.e. = 0.12x).

Second, we ran the LDAK software in a non-default mode in which every SNP has non-zero weight equal to the inverse of its LD score, so that two SNPs in perfect LD are assigned exactly the same weight instead of assigning one of the weights to zero (LDAK-nonzeroweights; see Methods and Table S5). (These weights attained likelihoods higher than the default LDAK model, though still much lower than the baseline-LD model (and the Gazal-LD model); see Figure S3.) We determined that LDAK enrichment estimates increased considerably (Figure S13a e.g. 5.28x, s.e. = 0.32x for conserved regions, 8.14x, s.e. = 1.40x for FANTOM5 enhancers^21^; higher than S-LDSC estimates for most annotations). (We also determined that the corresponding combined method, S-LDSC+LDAK-nonzeroweights (see Table S5), produces enrichment estimates nearly identical to S-LDSC+LDAK (*ρ_c_* = 0.99, no annotation significantly different; Figure S13b), confirming that the choice of LDAK weights does not impact our conclusions about S-LDSC.)

In summary, both of these experiments suggest that assigning zero heritability to a large majority of SNPs (or, more generally, an extremely imbalanced distribution of weights) may lead to downward bias in functional enrichment estimates.

## Discussion

Misspecification of heritability models can bias heritability and functional enrichment estimates. The S-LDSC approach of adding degrees of freedom to the model reduces the potential for model misspecification; in particular, the baseline-LD model^4^ infers the extent of MAF- and LD-dependent architectures directly from the data. Here, we observed that the baseline-LD model attains a higher likelihood than the LDAK model. We subsequently developed and evaluated a combined method (S-LDSC+LDAK) that flexibly incorporates the annotations used by both the S-LDSC and LDAK methods, serving as a “gold standard” for comparisons. In analyses of 16 UK Biobank traits, the results of S-LDSC+LDAK closely match the results of S-LDSC. We also note that an extension of S-LDSC with the baseline-LD model to low-frequency variants produces functional enrichment estimates for low-frequency variants (which are less impacted by LD) that are comparable to or larger than enrichment estimates for common variants^22^. Notably, we determined that lower enrichment estimates for the default version of LDAK could potentially be explained by the assignment of zero weights to most SNPs; this conclusion also applies to summary statistic based methods that assign zero weights to most SNPs^18^. We thus conclude that the S-LDSC method produces robust enrichment estimates (consistent with previous analyses of functional enrichment of fine-mapped causal SNPs; see Figure 2 of ref. ^23^ and Figure 3 of ref. ^24^, which reported large functional enrichments); we recommend using S-LDSC in preference to S-LDSC+LDAK in most settings, due to the complexities of computing LDAK model weights and running S-LDSC+LDAK.

Our results have several implications for future studies. First, they underscore the importance of accounting for LD-dependent architectures in analyses of functional enrichment, as suggested in recent work^3,4^; in particular, analyses using S-LDSC should be performed using the baseline-LD model^4^ instead of our earlier baseline model^5^, which produces slightly inflated estimates (see Supplementary Table 8c of ref. ^4^). Second, our original LD score regression (LDSC) method instead used the GCTA model to assess confounding in GWAS data^25^ and to estimate genetic correlations between traits^26^. We anticipate that S-LDSC with the baseline-LD model^4^ will improve upon results of LDSC with the GCTA model for these two applications. In particular, we determined that using S-LDSC with the baseline-LD model produced significant lower estimates of confounding than LDSC (regression intercept; Figure S14a), consistent with recent observations^18^; we thus recommend using S-LDSC with the baseline-LD model to assess confounding in GWAS data. Third, we previously suggested that, under strong model assumptions, LDSC could be used to estimate the heritability causally explained by all common SNPs^5,25,26^ (also see ref. ^27^); S-LDSC with the baseline-LD model may aid this endeavor, but further investigation is required. In particular, S-LDSC with the baseline-LD model produced higher heritability estimates than LDSC (Figure S14b), consistent with recent observations^18^. Fourth, we observed that the choice of SNP set can affect likelihood comparisons (see Figure 1a vs. Figure 1b). Thus, we recommend that future model comparisons should carefully consider the choice of SNP set, and should favor a SNP set that attains higher overall likelihoods (M = 4.6 million well-imputed HRC SNPs in our analyses; Figure 1b). Fifth, although we have included LDAK with non-zero weights (LDAK-nonzeroweights) only as a secondary analysis (because it differs from the recommended LDAK method^13^), this approach may warrant further investigation.

We note two limitations of the baseline-LD model. First, the baseline-LD model assumes that enrichment effects of all annotations combine additively, whereas the LDAK model assumes that LD and MAF effects combine multiplicatively; this motivates further work to assess whether LD and MAF effects, and functional annotation effects, may combine multiplicatively. Indeed, a recent study proposed a model that multiplies annotations from the LDAK and the baseline models^18^. However, we observed that this multiplicative model attained lower likelihoods than the baseline model when analyzing well-imputed HRC SNPs, and attained lower likelihood than the baseline-LD model when analyzing either well-imputed 1000G SNPs or well-imputed HRC SNPs (Figure S3). Second, the baseline-LD estimates 75 parameters, which may limit power in small data sets. However, the large data sets that we analyze here (*N* > 50K) allow S-LDSC to estimate these parameters with adequate power, as confirmed by the statistical significance of our model comparisons (Figure 1) and enrichment estimates (Figure 3 and Table S7).

Accurate estimation of components of heritability relies on accurate modeling of genetic architectures, and we anticipate that new models and corresponding methods will continue to improve our current knowledge. However, our results strongly support S-LDSC with the baseline-LD model for functional enrichment analyses.

## Methods

### The S-LDSC method

Stratified LD score regression (S-LDSC)^4,5^ is a method for partitioning heritability causally explained by common variants (minor allele frequency (MAF) ≥5%) across overlapping discrete or continuous annotations using genome-wide association study (GWAS) summary statistics and a linkage disequilibrium (LD) reference panel containing both common and low-frequency variants. The method rests on the idea that if an annotation *a* is associated to increased heritability, LD to variants with large values of *a* will increase the *χ*^2^ statistic of a variant more than LD to variants with small values of *a*. More precisely, S-LDSC models the vector *β* of per normalized genotype effect sizes as a mean-0 vector whose variance depends on *D* continuous-valued annotations *a*_1_,…, *a_D_*:

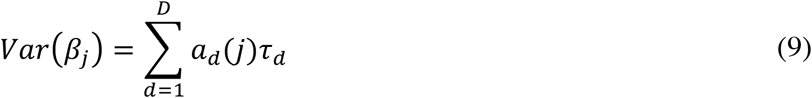

where *a_d_*(*j*) is the value of annotation *a_d_* at variant *j*, and *τ_d_* represents the per-variant contribution of one unit of the annotation *a_d_* to heritability. We can thus estimate the vector *τ* using the following relationship with the expected *χ*^2^ statistic of variant *j*:

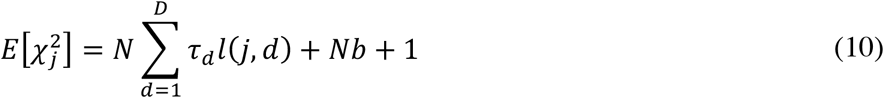

where 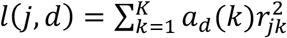 is the LD score of variant *j* with respect to continuous values *a_d_*(*k*) of annotation *a_d_*, *K* are the SNPs in a 1 cM window around *j, r_jk_* is the correlation between variant *j* and *k* in an LD reference panel, *N* is the sample size of the GWAS study, and *b* is a term that measures the contribution of confounding biases^25^. Then, the heritability causally explained by a subset of variants S can be estimated as 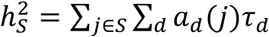.

Previous applications of S-LDSC^4,5^ used all SNPs with minor allele count ≥5 in unrelated whole genome sequenced individuals from the 1000 Genomes project^14^ as an LD reference panel to compute LD scores (this set of SNPs is denoted as the reference SNPs), restricted the regression between *χ*^2^ statistics and LD scores to estimate the *τ* parameters (Equation (10)) to the ~1M SNPs present in HapMap 3 as a proxy for well-imputed SNPs^5^ (this set of SNPs is denoted as the regression SNPs), and estimated heritability and enrichment values using the set of all reference SNPs with MAF ≥5% (this set of SNPs is denoted as the heritability SNPs). However, here we used UK10K^28^ as an LD reference panel (as in ref. ^22^), owing to its larger sample size (3,567 unrelated individuals) and closer match to the ancestry of UK Biobank samples. In our main S-LDSC analyses of UK Biobank data, we used all SNPs with minor allele count ≥5 as reference SNPs and estimated heritability and enrichment values using the set of all reference SNPs with MAF ≥5% (Table S4), to match previous applications of S-LDSC^4,5^. In all simulations, and in analyses of UK Biobank data using the LDAK estimand, we used all SNPs with MAF ≥1% as LD reference panel and estimated heritability and enrichment values using the set of all reference SNPs with MAF ≥1% (Table S4), to match LDAK^3^. We also ran constrained-intercept S-LDSC^26^ (which constrains the intercept, i.e. the *Nb* + 1 term from Equation (1), to equal 1) in the regression step when estimating *τ* values.

### The baseline-LD model

The original set of functional annotations used by S-LDSC is called the baseline model^5^, and contains *D* = 53 binary functional annotations including 28 main functional annotations (coding, conserved, DHS, histone marks, etc.) reflecting our knowledge of functional regions of the human genome^29,30^ at the time it was constructed (see Table S1). The model also contains 500bp windows around each binary annotation (and 100bp windows around 4 of the main annotations) to prevent enrichment estimates from being inflated by enriched heritability in flanking regions. More recently, we extended this baseline model to the baseline-LD model^4^, in order to take into account MAF- and LD-dependent architectures. The baseline-LD model contains *D* = 75 functional annotations including the 53 annotations from the baseline model, 6 new functional annotations, 10 MAF bins to account for MAF-dependent architectures, and 6 LD-related annotations (predicted MAF-adjusted allele age, level of LD in African populations, recombination rate, nucleotide diversity, a background selection statistic and CpG-content) to account for LD-dependent architectures (Table S1). (We note that we estimated that roughly half of LD-dependent causal effects were already captured by the functional annotations of the baseline model, as SNPs in functional annotations tend to have lower LD^4^.) The MAF bin boundaries were chosen so that each MAF bin contained the same proportion of reference variants, and 10 MAF bins were chosen because we determined that this was sufficient to produce robust results in simulations under different generative models^4^. The 6 LD-related annotations were included in the baseline-LD model because they all provide statistically significant signals of enriched heritability for real traits after conditioning on all other annotations in the baseline-LD model (see Figure 3c of ref. ^4^). In forward simulations, the 4 LD-related annotations that do not rely on empirical data were all conditionally informative for the selection coefficient of a variant, with effect sizes roughly proportional to the conditional effect sizes observed in heritability analyses of real traits, implying that the conditionally significant signals of enriched heritability are an expected consequence of negative selection^4^. We used baseline-LD version 1.1 (ref. ^31^) in all analyses (see URLs).

### The LDAK method

LDAK is a method for estimating heritability and functional enrichment from raw genotype-phenotype data^3^. When estimating the heritability explained by a single annotation *k*, the LDAK method uses the LDAK-annotation model:

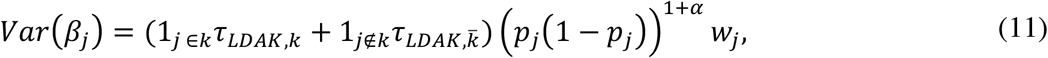

where 1_*j*∈*k*_ and 1_*j*∉*k*_ are indicator functions, *p_j_* is the allele frequency of SNP *j, α* is a parameter determining the relationship between MAF and heritability (recommended to be fixed at −0.25), and *w_j_* is the LDAK weight defined by minimizing the L1 or L2 norm of

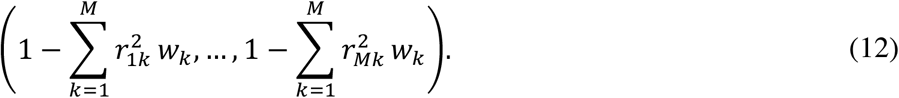

The coefficients *τ_LDAK,k_* and 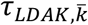 are estimated using restricted maximum likelihood^9,10^ (REML).

The LDAK method recommends that, for heritability partitioning analyses, users restrict their analyses to well-imputed SNPs (INFO score^16^ ≥0.99) with MAF ≥1% (ref. ^3^). We note that the LDAK software allows for incorporating INFO scores and could be extended to include poorly imputed SNPs. However, incorporating INFO scores has only been applied to heritability estimation with SNPs with INFO scores between 0.99 and 1.00, and has not been applied to heritability partitioning^3^.

### The S-LDSC+LDAK (S-LDSC estimand) and S-LDSC+LDAK (LDAK estimand) methods

We ran S-LDSC+LDAK (S-LDSC estimand) by using the S-LDSC method (with default options) on baseline-LD model annotations and two annotations constructed using LDAK weights (LDAK weights and allele frequencies were estimated using UK10K reference data).

We ran S-LDSC+LDAK (LDAK estimand) by restricting the LD reference panel to SNPs with MAF ≥1% in UK10K, partitioning the heritability of SNPs with MAF ≥1%, and estimating enrichment using the LDAK enrichment estimand (see Table S4 for the set of SNPs included in S-LDSC+LDAK enrichment estimands in our simulations and UK Biobank analyses).

### UK Biobank data and choice of independent traits

We analyzed data from the full UK Biobank release^7^ consisting of 487,409 samples genotyped on ~800K markers and imputed to ~93 million SNPs using the Haplotype Reference Consortium (HRC) dataset^15^ (*N* = 64,976 haplotypes from WGS data). We selected 16 quantitative traits that are heritable and independent, defined as having a heritability z-score > 7 with S-LDSC and pairwise phenotypic correlations below 0.1 (Table S2).

### Likelihood analyses

We restricted our likelihood analyses to *N* = 20K unrelated British-ancestry^32^ samples (due to LDAK computational constraints), who have phenotype information available for the 16 traits. We analyzed either genotypes imputed using the 1000 Genomes (1000G) reference panel^14^ (following imputation guidelines of ref. ^3^; *M* = 2.8 million well-imputed SNPs) or genotypes imputed using the Haplotype Reference Consortium (HRC) reference panel^15^ (M = 4.6 million well-imputed SNPs); in each case, we restricted our analyses to SNPs with INFO score^16^ ≥ 0.99 and MAF ≥ 1% (as in ref. ^3^), kept SNPs present in UK10K^28^, and excluded variants in the MHC region. We also replicated our analyses on *N* = 7K unrelated British-ancestry^32^ samples, using a sample size similar to ref. ^3^.

For all models, we computed allele frequency and/or LDAK weights in the *N* = 20K samples. For the baseline-LD and Gazal-LD models, we created 10 MAF bins from MAF ≥ 1% SNPs (compared to 5% in the default model), and recomputed the MAF-adjusted annotations and the nucleotide diversity annotation accordingly (see ref. ^4^). For the baseline-LD model, we removed the annotation containing all variants (leading to 74 total annotations), to remove co-linearity with the MAF bins (we note that the default baseline-LD model does not have this co-linearity issue, as the MAF bins span common SNPs and the annotation containing all variants span both common and low-frequency SNPs). The baseline-LD+LDAK model was constructed by adding to the baseline-LD annotations a continuous annotation with values 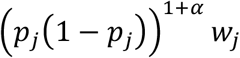.

We computed likelihoods using the LDAK software version 5.0 (see URLs). We used genotyping array (UK BiLEVE / UK Biobank), assessment center, sex, age and 20 principal components as fixed effect. We also included SNPs with P < 1 × 10^-20^ from single-SNP analysis (conditioned on the same covariates) and their correlated SNPs (correlation-squared > 0.5) as fixed effects, as in ref. ^3^. When computing likelihoods for the baseline-LD, Gazal-LD and baseline-LD+LDAK models, we estimated all *τ* parameters out-of-sample by running S-LDSC on mixed model association statistics computed using BOLT-LMM v2.3 (ref. ^19^) on the remaining unrelated British-ancestry UK Biobank samples, excluding the *N* = 20K samples and samples related to those samples (average *N* = 299K). S-LDSC was run by using the *N* = 20K samples as the LD reference panel, and using HapMap 3 SNPs as regression SNPs. We then computed values of per-SNP heritability for all well-imputed SNPs, and specified these values to the LDAK software as LDAK weights (while setting the *α* parameter to −1) to compute kinship matrixes. When computing likelihoods for the GCTA model, *α*-model and LDAK model, we did not estimate any parameters out-of-sample. All six models include one heritability parameter that is maximized in-sample when estimating the likelihood.

### Polygenic prediction analyses

We performed polygenic prediction using LDpred-funct-inf^17^ following the guidelines of Márquez-Luna et al.^17^; these analyses are analogous to the out-of-sample polygenic prediction approach proposed by ref. ^18^. We used association statistics computed by applying BOLT-LMM v2.3 (refs. ^19,33^) to *N* = 409K British-ancestry samples as training data, and *N* = 25K unrelated samples of other European ancestries (also unrelated to individuals from the training data) as validation data. We restricted our analyses to SNPs with minor allele frequency ≥ 1% and imputation accuracy ≥ 0.90, removed A/T and C/G SNPs, and kept SNPs present in UK10K (6,231,853 total SNPs). We used heritability estimates from BOLT-LMM as input to LDpred-funct-inf for all models. Per-SNP heritability for the *α*-model and the LDAK model were estimated using allele frequencies and LDAK weights computed using UK10K samples. Per-SNP heritability for the baseline-LD, Gazal-LD and baseline-LD+LDAK models were computed using *τ* parameters estimated out-of-sample by running S-LDSC with UK10K as the LD reference panel on the *N* = 409K British-ancestry samples.

### Simulations

We performed simulations to investigate the bias of S-LDSC, LDAK, and S-LDSC+LDAK under both S-LDSC and LDAK models. To overcome the issue of different S-LDSC and LDAK enrichment estimands (see above), we restricted our simulations to SNPs with MAF ≥1%, and modified S-LDSC to partition the heritability of SNPs with MAF ≥1% (see Table S4). We focused on the proportion of heritability explained by the annotations of interest rather than their enrichment, so that S-LDSC, LDAK and S-LDSC+LDAK estimate the same quantity (i.e. the proportion of heritability causally explained by SNPs with MAF ≥1%).

We performed simulations using chromosome 1 of the full UK Biobank release^7^ with imputed variants from the HRC. We restricted our simulations to 10,000 unrelated individuals, due to the computational limitations of the LDAK method. We sampled integer-valued genotypes from UK Biobank imputed dosages, restricting to 573,602 SNPs on chromosome 1 with MAF ≥1% and present in UK10K. We set trait heritability to *h*^2^ = 0.5 for all simulations and select *M* = 100,000 causal variants for main simulations (we repeated our simulations for different numbers of causal variants; Figure S6). We did not simulate genotype uncertainty as SNPs with an INFO score ≥ 0.99 (used by LDAK), and HapMap 3 SNPs (used by all methods with S-LDSC as the inference procedure) have very little genotype uncertainty.

We first simulated effect sizes using the baseline-LD model (including coding, conserved and DHS enrichment) with per-SNP heritability derived from the *τ* coefficients estimated across 31 independent traits^4^. More precisely, we computed 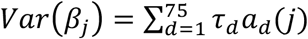 for each SNP, set negative values to 0, and simulated *β_j_* using a normal distribution with mean 0 and variance 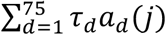. This produced MAF and LD effects similar to what was observed in ref. ^4^; for example, 20% of common SNPs with largest MAF (> 38%) explained 1.8x more heritability than the 20% with smallest MAF (< 10%), and the youngest 20% of common SNPs (based on MAF-adjusted predicted allele age) explained 3.9x more heritability than the oldest 20% (see Figure 4 of ref. ^4^ for other LD-related annotations). Second, we performed simulations under the LDAK-annotation model by selecting *M* = 100,000 causal variants and simulating effect sizes using a coding or conserved or DHS enriched architecture. Effect sizes were simulated using *α* = −0.25 (as recommended by LDAK authors^3^) and LDAK weights were computed using the 10,000 individuals. We simulated enrichment either by simulating larger effect sizes in the enriched annotation (and the same proportion of causal variants inside and outside the enriched annotation), or by simulating a larger proportion of causal variants in the enriched annotation (and same effect sizes for causal variants inside and outside the enriched annotation). We also performed simulation under the LDAK model, to test a model with no functional enrichment. For each simulation scenario, we performed 500 simulations.

S-LDSC was run on summary statistics generated for HapMap3 SNPs (95,335 chromosome 1 SNPs), and by using the baseline-LD model and UK10K^28^ (restricting to 589,319 UK10K SNPs of chromosome 1 with MAF ≥1%) as the LD reference panel (S-LDSC). To overcome the issue of unstable S-LDSC estimates under the LDAK generative model, we also ran S-LDSC with constrained intercept (S-LDSC0). We also ran S-LDSC with the baseline model^5^ (S-LDSC-baseline), and with the baseline model and constrained intercept (S-LDSC0-baseline), to assess the importance of modeling LD-dependent architectures. We ran 2 versions of LDAK for each simulation. First, we ran LDAK on all 573,602 SNPs (LDAK-allSNPs). Second, we ran LDAK on the subset of 361,050 well-imputed SNPs (INFO score^16^ ≥ 0.99) in all UK Biobank individuals (LDAK). We also ran S-LDSC+LDAK by adding the two LDAK coding-derived annotations (resp. conserved-derived or DHS-derived) to the baseline-LD model. To address the complexity of parameter estimation using the REML and S-LDSC procedure, we considered two auxiliary methods: LDAK-SS-allSNPs (see Table S5), a method that applies S-LDSC using the same LDAK-annotation model as the LDAK method, but the same set of SNPs (589,319 UK10K SNPs of chromosome 1 with MAF ≥ 1%) as the S-LDSC+LDAK (LDAK estimand) method; and LDAK-SS (see Table S5), a method that applies S-LDSC using the same LDAK-annotation model and the same set of SNPs as the LDAK method (361,050 UK10K SNPs of chromosome 1 with MAF ≥ 1% and INFO score ≥ 0.99). We note that the LDAK-SS method is similar to the method introduced in ref. ^18^. Results of LDAK-SS-allSNPs and LDAK-SS were compared to LDAK-allSNPs and LDAK, respectively.

In additional simulations under the baseline-LD+LDAK-annotation model (derived from the *τ* coefficients estimated across the 16 independent traits of this study), we also determined that S-LDSC+LDAK was unbiased, that S-LDSC slightly overestimated the proportion of heritability explained by conserved regions, and that LDAK was downwardly biased (Figure S5).

### Heritability partitioning analyses of UK Biobank data

For each of the 16 UK Biobank traits, we computed mixed model association statistics using BOLT-LMM v2.3 (ref. ^19^) with genotyping array (UK BiLEVE / UK Biobank), assessment center, sex, age, and age squared as covariates. We also included 20 principal components (included with the UK Biobank data release^7^) to correct for ancestry, as recommended by ref. ^19^. We included 672,292 directly genotyped SNPs in the mixed model (all autosomal biallelic SNPs with <10% missing data). S-LDSC and S-LDSC+LDAK were run using 3,567 whole genome sequenced UK10K^28^ samples (ALSPAC and TWINSUK cohorts) as the LD reference panel. S-LDSC and S-LDSC+LDAK (S-LDSC estimand) restricted their analyses to the 13,326,465 SNPs with minor allele count ≥5 (including 5,353,593 SNPs with MAF ≥5%). LDAK annotations for S-LDSC+LDAK (S-LDSC estimand) were constructed using UK10K allele frequencies and LDAK weights computed for all 13,326,465 UK10K SNPs with allele count ≥5 (note that LDAK software computes weight only for SNPs with MAF ≥1%). S-LDSC+LDAK (LDAK estimand) restricted its analyses to the 7,659,089 SNPs with MAF ≥1%. LDAK annotations for S-LDSC+LDAK (with LDAK estimand) were constructed using UK10K allele frequencies and LDAK weights computed for all 7,659,089 UK10K SNPs with MAF ≥1%. For each trait, S-LDSC was run once using the baseline-LD model version 1.1 (ref. ^31^), while S-LDSC+LDAK was run 28 times in turn for each of the functional annotations of interest.

LDAK was run as recommended by Speed et al.^3^. We considered *N* = 20K unrelated British-ancestry^32^ samples (due to the computational limitations of LDAK at larger sample sizes), and restricted our analyses to the 4,631,901 SNPs with MAF ≥ 1% and INFO score ≥ 0.99. We used *α* = −0.25 to compute kinship matrixes for each of the 28 functional annotations of interest. We ran restricted maximum likelihood (REML) using genotyping array (UK BiLEVE / UK Biobank), assessment center, sex, age and 20 principal components as fixed effect. We also included SNPs with P < 1 × 10^-20^ from single-SNP analysis (conditioned on the same covariates) and their correlated SNPs as fixed effects. To evaluate the impact of the proportion of SNPs with zero weights on functional enrichment estimates, we recomputed non-default LDAK weights as the inverse of the LD scores (LDAK-nonzeroweights), using LDAK options `--cut-weights sections --no-thin YES and --calc-weights sections --quick-weights YES`.

We partitioned the heritability of 16 UK Biobank traits across 28 main functional annotations from the baseline model using different methods. We meta-analyzed enrichments using random-effects meta-analyses implemented in the R package *rmeta*. We performed pairwise comparisons of methods using the concordance correlation coefficient^20^ (*ρ_c_*) defined as

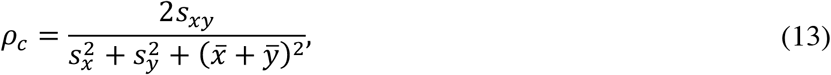

where *s_xy_* is the covariance of estimates of methods *x* and *y*, 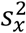 and 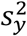 are the variances for estimates of methods *x* and *y*, respectively, and 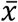 and 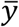 are the means for estimates of methods *x* and *y*, respectively. We tested whether two different enrichment estimates for the same annotation were significantly different by computing a Z score based on the respective estimates and standard errors. We note that it is possible for errors to be correlated, as they are computed using the same data; if this is the case, our test for significantly different enrichment estimates is conservative.

## URLs

S-LDSC, https://github.com/bulik/ldsc. Baseline-LD model version 1.1, https://data.broadinstitute.org/alkesgroup/LDSCORE/1000G_Phase3_baselineLD_v1.1_ldscores.tgz. LDAK v5, http://dougspeed.com/downloads/. BOLT-LMM v2.3, https://data.broadinstitute.org/alkesgroup/BOLT-LMM/.

## Code and data availability

S-LDSC is available at https://github.com/bulik/ldsc. Baseline-LD model version 1.1 annotations and LD scores are available at https://data.broadinstitute.org/alkesgroup/LDSCORE/1000G_Phase3_baselineLD_v1.1_ldscores.tgz. LDAK version 5 is available at http://dougspeed.com/downloads/. UK Biobank association statistics, computed using BOLT-LMM v2.3, are available at http://data.broadinstitute.org/alkesgroup/UKBB/.

## Acknowledgements

We are grateful to Po-Ru Loh for assistance with UK Biobank data and to Luke O’ Connor, Doug Speed and David Balding for helpful discussions. This research was conducted using the UK Biobank Resource under Application 16549 and was supported by NIH grants U01 HG009379, R01 MH101244 and R01 MH107649.

## Supplementary Note

### Consistent estimands are essential when comparing results for LD-related annotations

S-LDSC and LDAK use different functional enrichment estimands. We determined that the different estimands make little difference in analyses of functional annotations, as S-LDSC+LDAK (S-LDSC estimand) and S-LDSC+LDAK (LDAK estimand) produce similar enrichment estimates (Figure S10). However, consistent estimands are essential when comparing results for annotations related to LD or MAF.

We consider a “thinned SNPs” annotation, constructed as in ref. ^34^ such that one of each pair of SNPs within 1cM with *r*^2^>0.2 are removed. This annotation is enriched in SNPs with low LD that cannot be tagged by other SNPs. We constructed a thinned annotation from all UK10K SNPs with an allele count ≥ 5 for estimation using S-LDSC and S-LDSC+LDAK (S-LDSC estimand), from all UK10K SNPs with MAF ≥ 1% for estimation using S-LDSC+LDAK (LDAK estimand), and from all UK Biobank SNPs with MAF ≥ 1% and INFO score ≥ 0.99 for estimation using LDAK. These annotations contain 2.16%-4.85% of SNPs and are expected to explain between 23.71%-36.10% of heritability under the LDAK model (Table S8). We observed that all methods estimated high enrichment when using the proportion of SNPs as the denominator (S-LDCS estimand; 3.38x-3.87x enrichment), but depleted enrichment when using the expected proportion of heritability under the LDAK model as the denominator (LDAK estimand; 0.31x-0.55x enrichment) (Table S8). In particular, comparing S-LDSC (enrichment = 3.87x, s.e. = 0.40x) and LDAK (enrichment = 0.45x, s.e. = 0.05x) without accounting for the different estimands would lead to incorrect conclusions.

We provide one additional example to illustrate the difference between the S-LDSC estimand and the LDAK estimand. Consider the set of all SNPs with MAF≥20%. The baseline-LD model used by S-LDSC and the LDAK model are in agreement that these SNPs have higher per-SNP heritability than other SNPs. Using the S-LDSC estimand (denominator = proportion of SNPs), SNPs with MAF≥20% are enriched for heritability. Using the LDAK estimand (denominator = expected proportion of heritability under the LDAK model), SNPs with MAF≥20% are not enriched for heritability. Neither estimand is correct or incorrect, but comparing results of S-LDSC and LDAK without accounting for the different estimands would lead to incorrect conclusions.

### Unstable S-LDSC estimates in simulations under the LDAK model

In our simulations under the LDAK and LDAK-annotation models, we observed unstable estimates for S-LDSC, but not for S-LDSC with constrained intercept. We hypothesized that this result is due to the relatively flat relationship between LD scores and *χ*^2^ statistics in simulations under the LDAK model (Figure S1), as compared to what is observed in real data (Figure S15) or in simulations under the baseline-LD model (Figure S1). Indeed, in these simulations, S-LDSC tends to interpret the inflated *χ*^2^ statistics (relative to a null distribution) as a consequence of confounding (by estimating high intercept and a slope close to 0) rather than due to heritability; constraining the intercept to 1 forces S-LDSC to interpret the inflated *χ*^2^ statistics as a consequence of heritability (Figure S16).

### Differences between S-LDSC and LDAK are not due to inference methods, sample sizes, or sets of SNPs

We hypothesized that the large difference between LDAK and S-LDSC+LDAK (LDAK estimand) enrichment estimates for conserved regions (Figure 3b) could be due to misspecification of the LDAK model; other potential explanations include different inference methods (S-LDSC vs. REML), different sample sizes (average *N* = 434K vs. *N* = 20K), and/or different sets of SNPs (7.7M SNPs with MAF ≥ 1% vs. 4.6M SNPs with MAF ≥ 1% and INFO score ≥ 0.99).

To investigate these alternative explanations, we compared LDAK to three auxiliary methods, which infer the parameters of the LDAK-annotation model or the baseline-LD model using S-LDSC, and require only summary statistics and an LD reference panel. These auxiliary methods are summarized in Table S5.

To investigate the impact of the different inference methods and different sample sizes, we compared LDAK to LDAK-SS, a method that applies S-LDSC with average *N* = 434K using the same model (LDAK-annotation) and set of SNPs (4.6M SNPs with MAF ≥ 1% and INFO score ≥ 0.99) as the LDAK method (see Table S5). We note that the LDAK-SS method is similar to the method introduced in ref. ^18^. In our simulations, we observed that LDAK-SS produced results similar to LDAK (Figure S5), demonstrating that the S-LDSC inference method can accurately estimate functional enrichment under the LDAK model. In the analysis of 16 UK Biobank traits, we observed similar enrichment estimates (*ρ_c_* = 0.91, no annotation significantly different, LDAK-SS conserved enrichment = 2.44x, s.e. = 0.08x versus 2.60x, s.e. = 0.17x for LDAK; Figure S17), indicating that the lower enrichment estimates of LDAK (and LDAK-SS) as compared to S-LDSC+LDAK (LDAK estimand) cannot be explained by differences in inference method or sample size.

To investigate the impact of the different sets of SNPs, we performed two comparisons. First, we compared LDAK to LDAK-SS-allSNPs, a method that applies S-LDSC (with average *N* = 434K) using the same model (LDAK-annotation) as the LDAK method, but the same set of SNPs (7.7M SNPs with MAF ≥ 1%) as the S-LDSC+LDAK (LDAK estimand) method (see Table S5). In our simulations, we observed that LDAK-SS-allSNPs produced results nearly identical to LDAK-allSNPs (Figure S5). In the analysis of 16 UK Biobank traits, we again observed similar enrichment estimates (*ρ_c_* = 0.81, 3 annotations significantly different, LDAK-SS-allSNPs conserved enrichment = 3.13x, s.e. = 0.13x; Figure S18), indicating that the low LDAK conserved enrichment cannot be explained by restricting to well-imputed SNPs. Next, we compared LDAK to S-LDSC-LDAKSNPs, a method that applies S-LDSC (with average *N* = 434K) using the baseline-LD model, analyzing the same set of SNPs as the LDAK method (4.6M SNPs with MAF ≥ 1% and INFO score ≥ 0.99) (see Table S5). We observed less concordant results and a high conserved enrichment (*ρ_c_* = 0.49, 6 annotations significantly different, S-LDSC-LDAKSNPs conserved enrichment = 6.77x, s.e. = 0.38x; Figure S19), indicating that the high conserved enrichment of S-LDSC+LDAK (LDAK estimand) is not a consequence of the inclusion of poorly imputed SNPs.

Having ruled out inference method, sample size, and SNP set as potential explanations for the difference between LDAK and S-LDSC+LDAK (LDAK estimand), we thus concluded that the low LDAK conserved enrichment estimates is likely due to LDAK model misspecification, consistent with the lower LDAK likelihood in formal model comparisons (Figure 1) and strong downward bias in LDAK enrichment estimates in simulations under the baseline-LD model (Figure S5; also see Figure 2).

See Excel file.

**Table S1: Functional annotations of the baseline-LD model.** We list the 75 annotations of the baseline-LD model, with accompanying information. The “Main annotation” column indicates whether the annotation is included as one of the 28 main functional annotations.

See Excel file.

**Table S2: UK Biobank independent traits.** We list the 16 UK Biobank traits analyzed, with accompanying information.

See Excel file.

**Table S3: Likelihood of different models of per-SNP heritability.** We report the log likelihood of nine different per-SNP heritability models over 16 independent UK Biobank quantitative traits. (**a**) Datasets using *N* = 7K and *M* = 2,835,699 well-imputed 1000G SNPs (as in ref. ^3^). (**b**) Datasets using *N* = 7K and *M* = 4,631,901 well-imputed HRC SNPs. (**c**) Datasets using *N* = 20K and *M* = 2,835,699 well-imputed 1000G SNPs (as in ref. ^3^). (**d**) Datasets using *N* = 20K and *M* = 4,631,901 well-imputed HRC SNPs.

See Excel file.

**Table S4: Reference SNPs, Regression SNPs and Heritability SNPs used by S-LDSC, LDAK and S-LDSC+LDAK.** For S-LDSC and S-LDSC+LDAK we list the Reference SNPs (used to compute LD scores), Regression SNPs (included in the regression) and Heritability SNPs (used to compute proportion of heritability and enrichment). For LDAK, we list the Heritability SNPs (used to run LDAK and compute proportion of heritability and enrichment), as References SNPs and Regression SNPs are not applicable. We provide this information for both Simulations (column 1) and Analyses of 16 UK Biobank traits (column 2). In the simulations, we restricted to causal SNPs with MAF ≥ 1%, and adapted the S-LDSC and S-LDSC+LDAK Reference SNPs and Heritability SNPs to match LDAK (i.e. SNPs with MAF ≥ 1%). In the analyses of 16 UK Biobank traits, we ran S-LDSC and LDAK using the recommended settings^4^, and ran two versions of S-LDSC+LDAK using S-LDSC and LDAK estimands respectively.

**Table S5:** Methods for estimating functional enrichment. We summarize the four main methods and the seven auxiliary methods for estimating functional enrichment. For each method, we list their input data, per-SNP heritability model, inference method, estimand and SNP set (either the set of reference SNPs used to compute LD scores for methods using the S-LDSC inference method, or the set of SNPs included in the GRM for methods using the REML inference method). Methods that analyze summary statistics (SS) were applied to summary statistics computed using *N* = 434K individuals; methods that analyze raw genotype/phenotype data (Raw) were applied to *N* = 20K individuals (see main text). The LDAK enrichment estimand for an annotation is defined as the proportion of heritability causally explained by genotyped SNPs in the annotation, divided by the expected proportion of heritability under the LDAK model, in a model in which only SNPs with MAF ≥ 1% are causal. The S-LDSC enrichment estimand for an annotation is defined as the proportion of heritability causally explained by SNPs in the annotation with MAF ≥ 5% from the LD reference panel, divided by the proportion of SNPs with MAF ≥ 5% in the LD reference panel that fall in the annotation. LDAK-allSNPs was applied only in simulations and not in our application on the genotype imputed data of UK Biobank traits, as LDAK authors recommend to apply their method on well-imputed SNPs only.

| Method | Data | Model | Inference | Estimand | SNP set | Ref |
| --- | --- | --- | --- | --- | --- | --- |
| *Main methods* |  |  |  |  |  |  |
| S-LDSC | SS | Baseline-LD | S-LDSC | S-LDSC | AC $\geq 5$<br>($M = 13.3\text{M}$ UK10K SNPs) | <sup>4</sup> |
| LDAK | Raw | LDAK-annotation | REML | LDAK | MAF $\geq 1\%$, INFO $\geq 0.99$<br>($M = 4.6\text{M}$ HRC SNPs) | <sup>3</sup> |
| S-LDSC+LDAK<br>(S-LDSC estimand) | SS | Baseline-LD+<br>LDAK-annotation | S-LDSC | S-LDSC | AC $\geq 5$<br>($M = 13.3\text{M}$ UK10K SNPs) | - |
| S-LDSC+LDAK<br>(LDAK estimand) | SS | Baseline-LD+<br>LDAK-annotation | S-LDSC | LDAK | MAF $\geq 1\%$<br>($M = 7.7\text{M}$ UK10K SNPs) | - |
| *Auxiliary methods* |  |  |  |  |  |  |
| LDAK-SS | SS | LDAK-annotation | S-LDSC | LDAK | MAF $\geq 1\%$, INFO $\geq 0.99$<br>($M = 4.6\text{M}$ UK10K SNPs) | - |
| LDAK-SS-allSNPs | SS | LDAK-annotation | S-LDSC | LDAK | MAF $\geq 1\%$<br>($M = 7.7\text{M}$ UK10K SNPs) | - |
| S-LDSC-LDAKSNPs | SS | Baseline-LD | S-LDSC | LDAK | MAF $\geq 1\%$, INFO $\geq 0.99$<br>($M = 4.6\text{M}$ UK10K SNPs) | - |
| S-LDSC-nonzeroLDAKSNPs | SS | Baseline-LD | S-LDSC | LDAK | MAF $\geq 1\%$, INFO $\geq 0.99$,<br>LDAK weight $\neq 0$<br>($M = 708\text{K}$ UK10K SNPs) | - |
| LDAK-nonzeroweights | Raw | LDAK-annotation<br>(non-zero weights) | REML | LDAK | MAF $\geq 1\%$, INFO $\geq 0.99$<br>($M = 4.6\text{M}$ HRC SNPs) | - |
| S-LDSC+LDAK-<br>nonzeroweights<br>(LDAK estimand) | SS | Baseline-LD+<br>LDAK-annotation<br>(non-zero weights) | S-LDSC | LDAK | MAF $\geq 1\%$<br>($M = 7.7\text{M}$ UK10K SNPs) | - |
| LDAK-allSNPs* | Raw | LDAK-annotation | REML | LDAK | MAF $\geq 1\%$ | - |

SS: summary statistics; AC: allele count; INFO: INFO score; \*: applied only in simulations.

See Excel file.

**Table S6: Simulation results.** We report numerical results from **Figure S5.**

See Excel file.

**Table S7: UK Biobank functional enrichment estimates.** We report enrichment estimates of the 28 main functional annotations, estimated across 16 independent UK Biobank traits. For each annotation, we report the proportion of SNPs of the annotation. For methods using the LDAK estimand, we report the proportion of heritability expected under the LDAK model.

See Excel file.

**Table S8: Enrichment for a thinned SNPs annotation.** We report results of S-LDSC (using baseline-LD model annotations and a thinned SNPs annotation), S-LDSC+LDAK (S-LDSC estimand) (using baseline-LD model annotations, a thinned SNPs annotation, and 2 thinned SNPs annotations constructed from LDAK weights), S-LDSC+LDAK (LDAK estimand) (using baseline-LD model annotations, a thinned SNPs annotation, and 2 thinned SNPs annotations constructed from LDAK weights), and LDAK (using 2 thinned SNPs annotations constructed from LDAK weights). The thinned SNPs annotations were built from all UK10K SNPs with an allele count greater than 5 for S-LDSC (with the baseline-LD model annotations and the thinned SNP annotation) and the S-LDSC+LDAK (S-LDSC estimand), from all UK10K SNPs with a MAF ≥ 1% for S-LDSC+LDAK (LDAK estimand), and from all UK Biobank SNPs with a MAF ≥ 1% and a INFO score ≥ 0.99 for LDAK. We put in grey the denominator and the enrichment that does not correspond to appropriate estimand.

**Figure S1:**
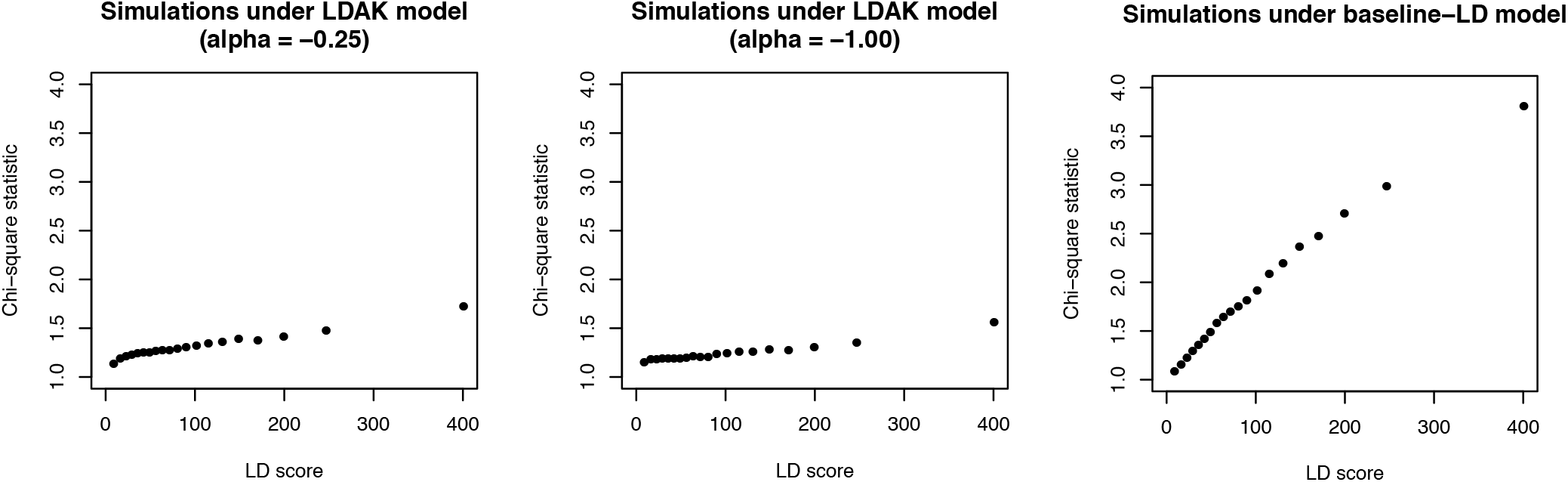
Relationship between LD scores and *χ*^2^ statistics in simulations under the LDAK and baseline-LD models. For each of 20 LD score bins, we report the mean LD score (computed on UK10K) and the mean *χ*^2^ statistics computed in 20 simulations under LDAK model (*α* = −0.25) (left panel), LDAK model without MAF-dependent architecture (*α* = −1.00) (middle panel), and baseline-LD model (right panel). (The analogous plots for real traits are provided in Figure S15.) Results are restricted to HapMap 3 regression SNPs used by S-LDSC to perform the regression (results were similar when including all UK10K SNPs; data not shown). Simulations under the LDAK model (left panel) show a relatively flat relationship between LD scores and *χ*^2^ statistics. Simulations under the LDAK model without MAF-dependent architecture (middle panel) show an even flatter relationship between LD scores and *χ*^2^ statistics, as the MAF-dependent architecture tends to increase *χ*^2^ statistics of SNPs with high LD (since common SNPs have higher LD). Simulations under the baseline-LD model (right panel) show a strong relationship between LD scores and *χ*^2^ statistics. We note that, according to the authors of LDAK, *“Our aim is that the variance terms are equal, which we can achieve by solving a matrix equation of the form Cw =* 1” (ref. ^1^, Appendix A), where *C* is a matrix of squared correlations between SNPs and *w* is a vector of LDAK weights. This aim is precisely the outcome that the variance of marginal association statistics is independent of LD. (This implies that recent population bottlenecks and founder events (which increase LD) should greatly reduce causal effect sizes.) In simulations under the LDAK model (left panel), we observed a relationship between LD scores and *χ*^2^ statistics that is not perfectly flat, such that the aim was not achieved; we hypothesize that this may be because the minor allele frequency term in the updated LDAK model (Equation (1) of ref. ^3^) partially counteracts the effects of their weights and/or because the outcome *Cw* = 1 is difficult to achieve. In simulations under the LDAK model without MAF-dependent architecture (middle panel), we also observed a relationship between LD scores and *χ*^2^ statistics that is not perfectly flat; we hypothesize that this may be because the outcome *Cw* = 1 is difficult to achieve. We finally note that the validity of the LDAK model depends on the LD-dependence of *causal* effect sizes, and is not an automatic consequence of tagging; indeed, many studies have shown that methods that do not apply LD-dependent weights obtain unbiased estimates in simulations that include LD but do not include LD-dependence of *causal* effect sizes^3,9^.

**Figure S2:**
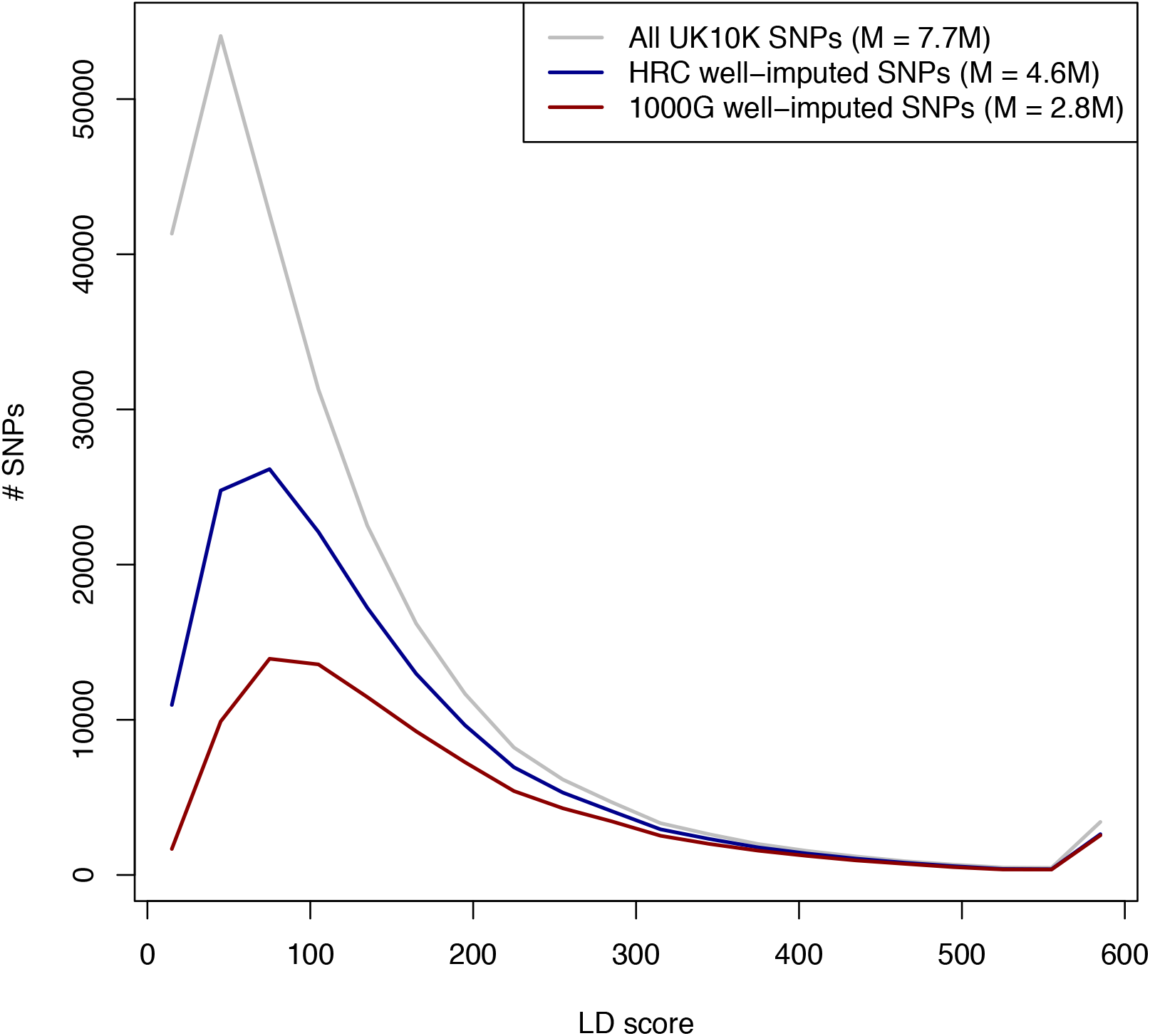
Comparison of LD scores in different datasets. We computed LD scores on 7.7M SNPs with MAF ≥ 1% in UK10K. We plotted repartition of those LD scores for all SNPs (median LD score = 82), 4.6M well-imputed HRC SNPs (median LD score = 110), and 2.8M well-imputed 1000G SNPs (median LD score = 139).

**Figure S3A:**
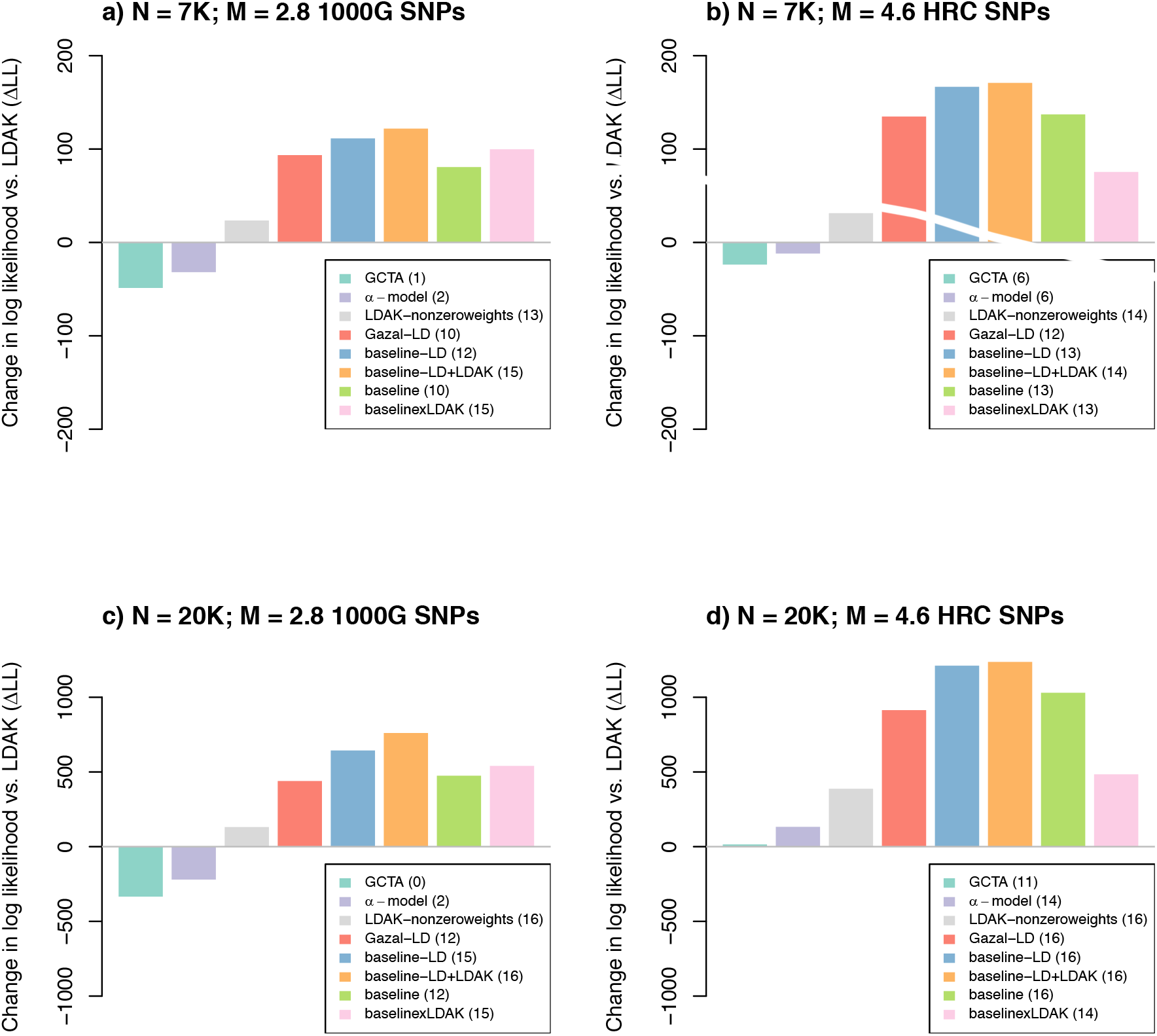
Likelihood comparison of different models of per-SNP heritability. We report the change in log likelihood compared to the LDAK model (ΔLL) of eight other per-SNP heritability models, summed across 16 independent UK Biobank quantitative traits. All models include one heritability parameter that is maximized insample when estimating the likelihood. (**a**) Analyses using *N* = 7K samples (as in ref. ^3^) and *M* = 2,835,699 well-imputed 1000G SNPs (as in ref. ^3^). (**b**) Analyses using *N* = 7K samples (as in ref. ^3^) and *M* = 4,631,901 well-imputed HRC SNPs. (**c**) Analyses using *N* = 20K samples and *M* = 2,835,699 well-imputed 1000G SNPs (as in ref. ^3^). (**d**) Analyses using *N* = 20K samples and *M* = 4,631,901 well-imputed HRC SNPs. Numbers between parentheses in figure legends indicate the number of traits with ΔLL > 0. Numerical results, including results for each trait, are reported Table S3. For comparison purposes, we also report likelihood results for the LDAK-nonzeroweights, a non-default version of LDAK that models SNPs in perfect LD differently by assigning nonzero weights to all SNPs. We also report likelihood results for the baseline model^5^ (whose enrichment estimates using S-LDSC were compared to LDAK enrichment estimates in ref. ^3^). We also report likelihood results for a model that multiplies annotations from the baseline and LDAK models (baselinexLDAK; similar to the model introduced in ref. ^18^), in which 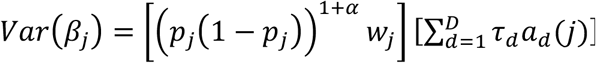, with *D* represents the functional annotations of the baseline-LD model (i.e. all annotations except the 10 MAF bins and the 6 LD-related annotations). This model attained lower likelihoods than the baseline-LD model (for 8, 11, 12, and 16 traits in panels a, b, c, and d, respectively), showing that adding MAF and LD annotations to *Var*(*β_j_*) is more informative than multiplying *Var*(*β_j_*) by (*p_j_*(1 – *p_j_*))^1+*α*^ *w_j_*. We also determined that the baselinexLDAK model attains a higher likelihood than the baseline model only when applied to datasets using well-imputed 1000G SNPs, but not when using well-imputed HRC SNPs. Numerical results, including results for each trait, are reported in Table S3.

**Figure S3B:**
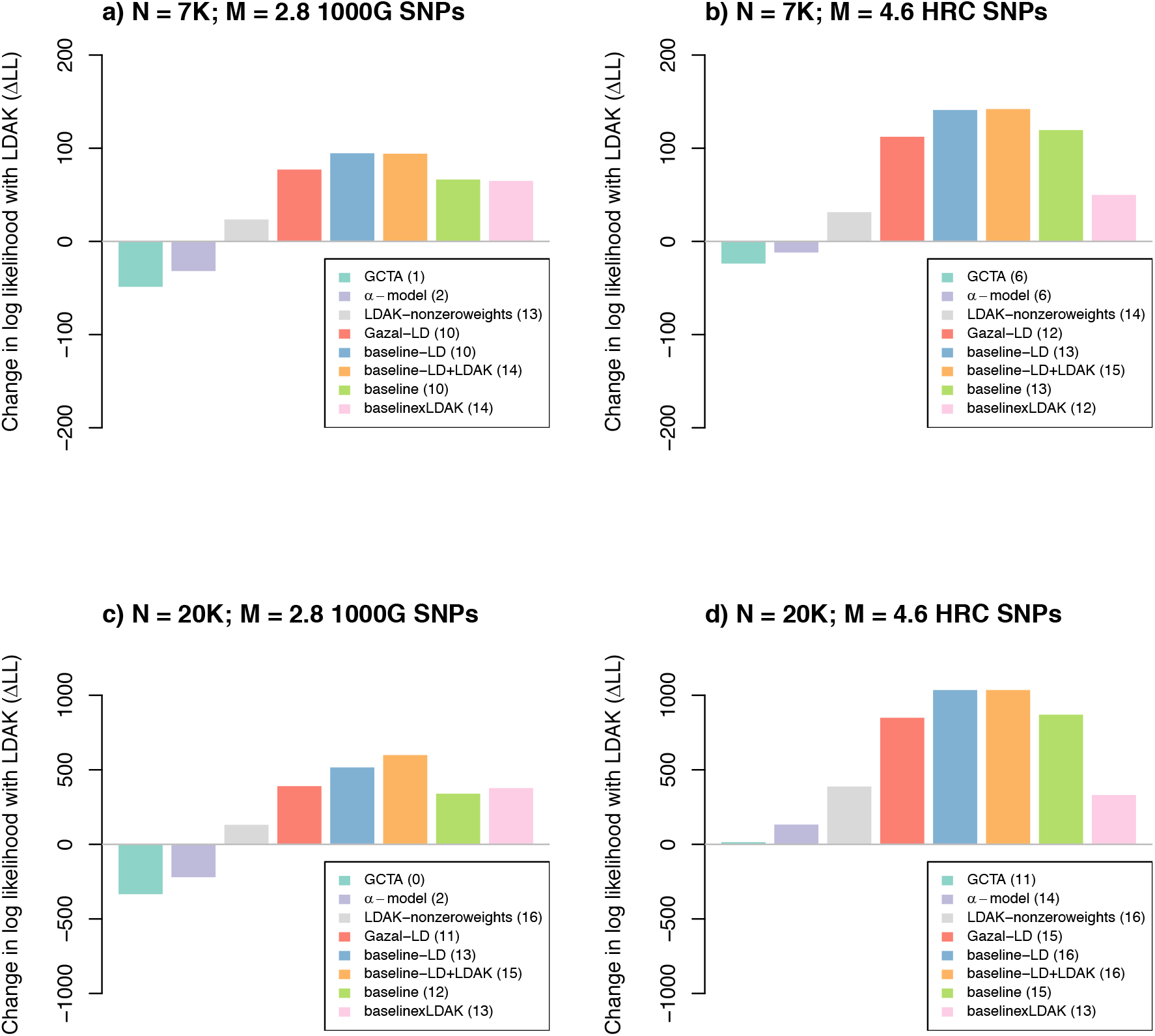
Likelihood comparison of different models of per-SNP heritability where we trained model parameters using a meta-analysis across 16 traits. We report the change in log likelihood compared to the LDAK model (ΔLL) of eight other per-SNP heritability models, summed across 16 independent UK Biobank quantitative traits. All models include one heritability parameter that is maximized in-sample when estimating the likelihood. To compute per-SNP heritability of the Gazal-LD, baseline-LD, baseline-LD+LDAK, baseline and baselinexLDAK models, we meta-analyzed the *τ* parameters estimated out-of-sample across the 16 traits, so that each model uses per-SNP heritability that is not trait-specific (as compared in Figure S3A). (**a**) Analyses using *N* = 7K samples (as in ref. ^3^) and *M* = 2,835,699 well-imputed 1000G SNPs (as in ref. ^3^). (**b**) Analyses using *N* = 7K samples (as in ref. ^3^) and *M* = 4,631,901 well-imputed HRC SNPs. (**c**) Analyses using *N* = 20K samples and *M* = 2,835,699 well-imputed 1000G SNPs (as in ref. ^3^). (**d**) Analyses using *N* = 20K samples and *M* = 4,631,901 well-imputed HRC SNPs. We observed similar results than in Figure S3A.

**Figure S4:**
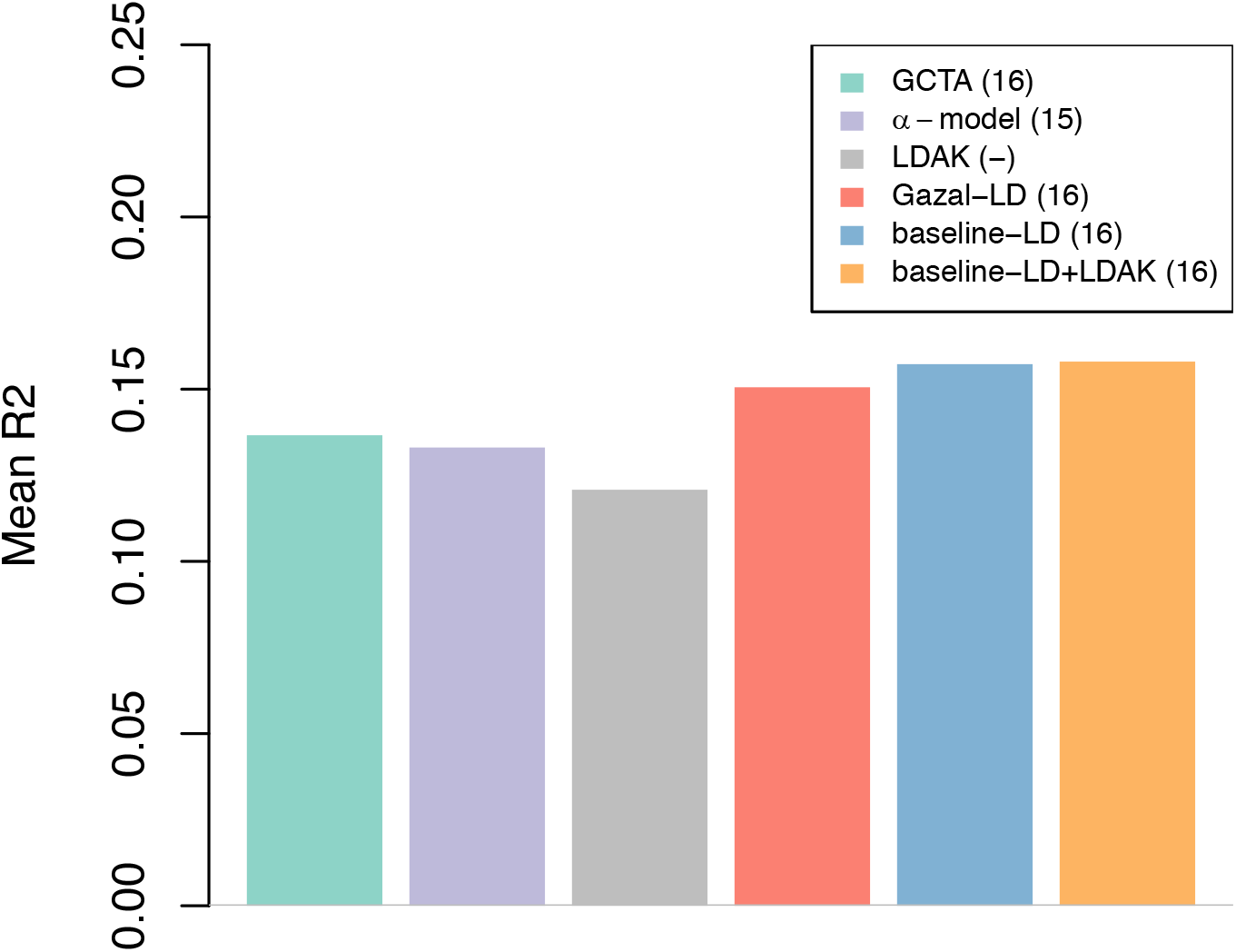
Out-of-sample polygenic prediction of different models of per-SNP heritability. We report the mean squared correlation (R2) between true phenotypes and predicted phenotypes using LDpred-funct-inf^17^ with six different per-SNP heritability models, across 16 independent UK Biobank traits (*N* = 409K training samples of British ancestry, *N* = 25K validation samples of other European ancestries, *M* = 6,645,003 UK10K SNPs). Numbers between parentheses in the figure legend indicate the number of traits where R2 is higher than LDpred-funct-inf with the LDAK model.

**Figure S5:**
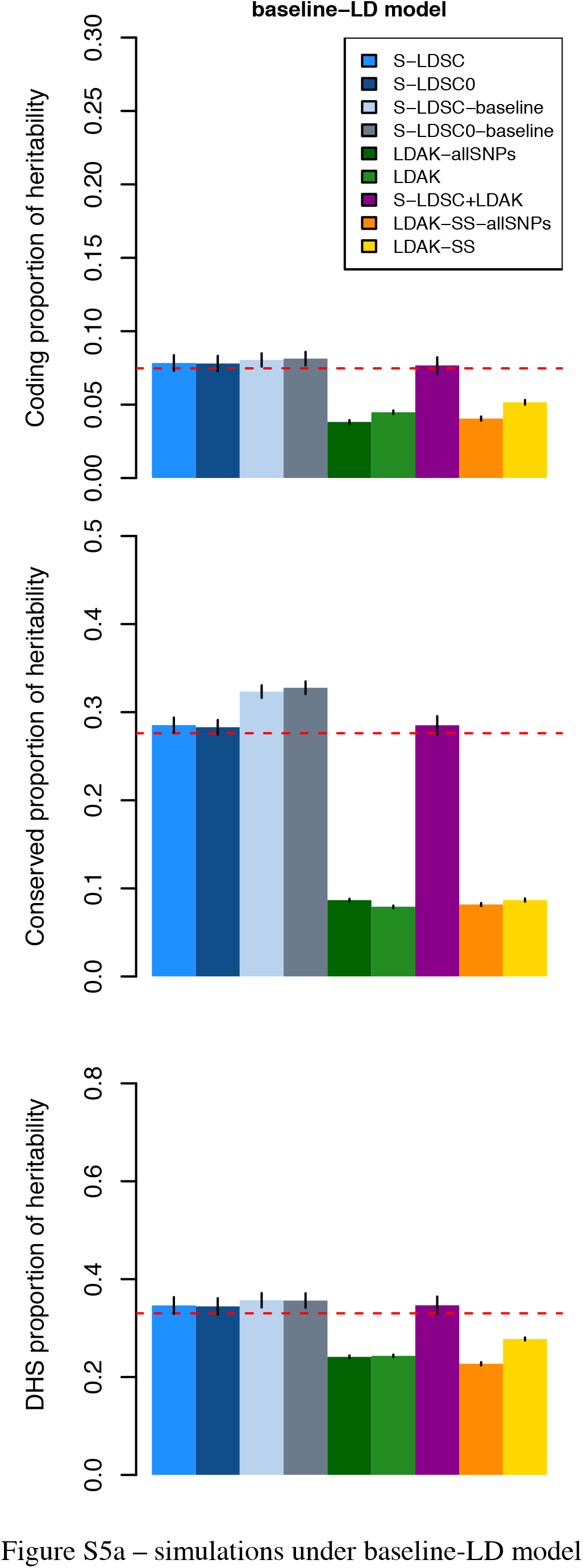
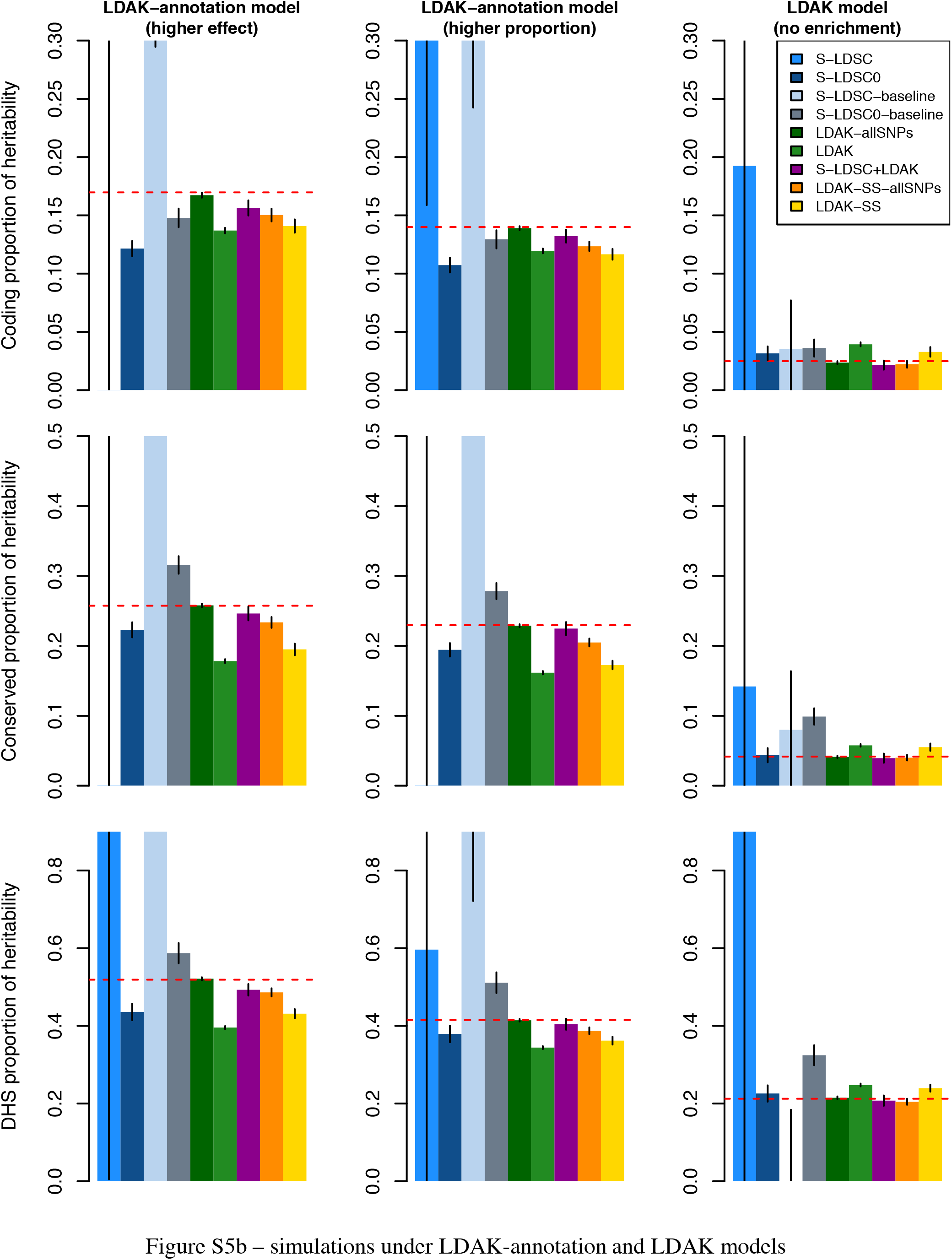
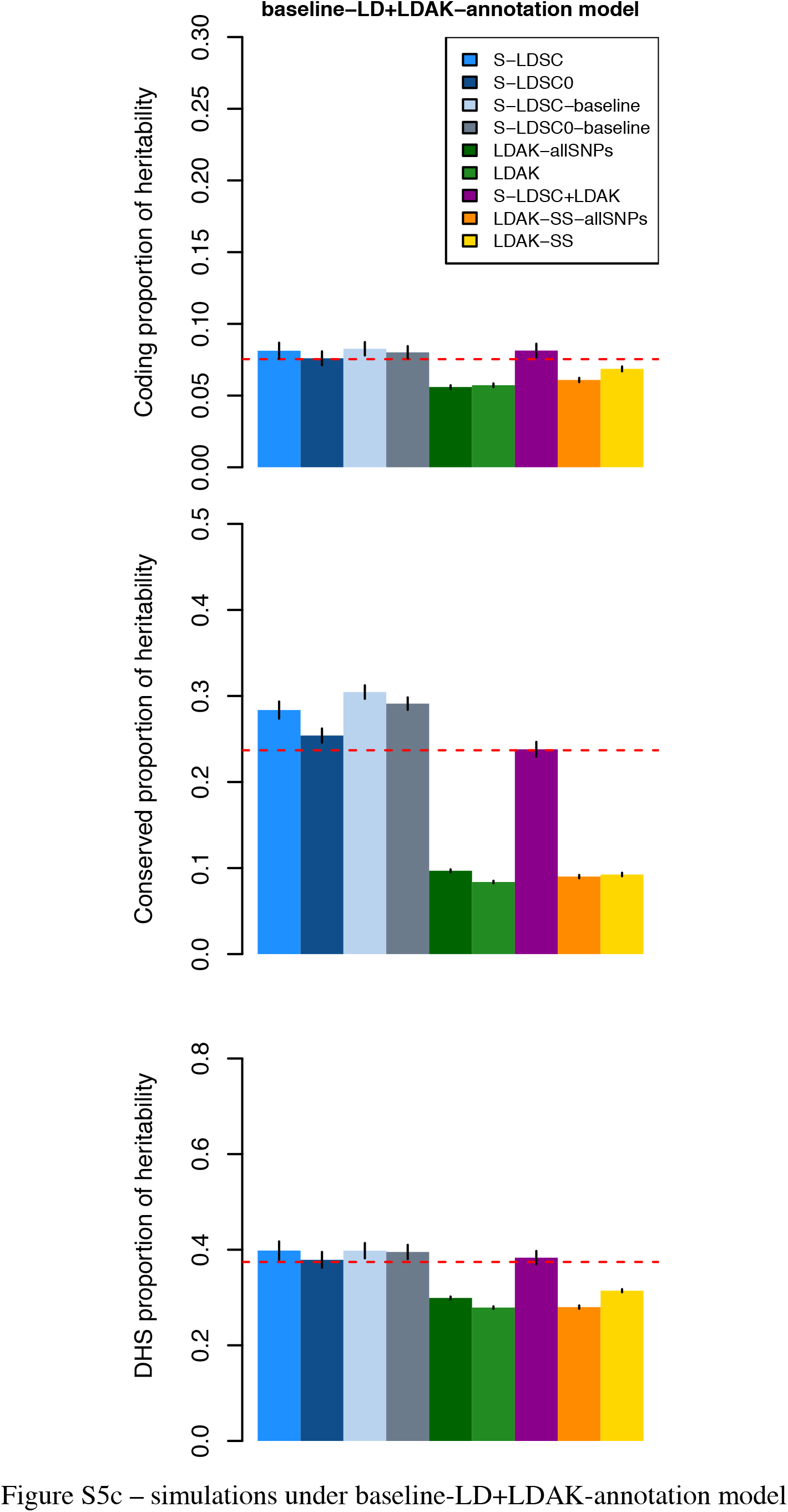
Simulations to assess LDAK, S-LDSC and S-LDSC+LDAK enrichment accuracy. We report enrichment simulations of 9 different methods, 3 different enriched annotations, and 5 different simulation scenarios involving 4 different generative models. The 9 different methods are S-LDSC with the baseline-LD model (S-LDSC), S-LDSC with the baseline-LD model and with intercept constrained to 1 (S-LDSC0), S-LDSC with the baseline model (S-LDSC-baseline), S-LDSC with the baseline model and with intercept constrained to 1 (S-LDSC0-baseline), LDAK using all SNPs (LDAK-allSNPs), LDAK with default SNP filtering, i.e. restricted to SNPs with INFO score ≥0.99 (LDAK), S-LDSC with the baseline-LD model and two annotations based on LDAK weights (S-LDSC+LDAK), S-LDSC with two annotations based on LDAK weights (LDAK-SS-allSNPs), and S-LDSC with two annotations based on LDAK weights and restricted to the same SNPs than the LDAK method (LDAK-SS). The 3 different enriched annotations are coding (row 1), conserved (row 2), and DHS (row 3). The 4 different generative models are baseline-LD (using the baseline-LD model with previously estimated parameters^4^) (**a**), LDAK-annotation and LDAK (**b**), and baseline-LD+LDAK-annotation (using the baseline-LD+LDAK-annotation model with parameters estimated in this study) (**c**). The 3 different simulation scenarios in panel (**b**) are functional enrichment with same proportion of causal SNPs in the enriched annotation but higher effect size variance (column 1), functional enrichment with higher proportion of causal SNPs in the enriched annotation but same effect size variance (column 2), and no enrichment (column 3). Panel (**a**) shows that in simulations under the baseline-LD model, 1) the four S-LDSC methods and S-LDSC+LDAK method are unbiased, and 2) other LDAK-related methods are downward biased. Panel (**b**) shows that, 1) the LDAK-allSNPs is unbiased, 2) S-LDSC+LDAK is slightly downward biased or unbiased, 3) LDAK is downward biased when simulating functional enrichment and upward biased under no enrichment, 4) S-LDSC produces unstable estimates while S-LDSC0 produces a stable downward bias, and 5) S-LDSC-baseline produces unstable estimates while S-LDSC0-baseline produces a stable upward bias (as in ref. ^18^). Panel (**c**) confirms that in simulations under the S-LDSC+LDAK-annotation model that S-LDSC+LDAK is unbiased. Dashed red lines indicate true values. Error bars represent the 95% confidence intervals based on 500 simulations. See Table S6 for numerical results.

**Figure S6:**
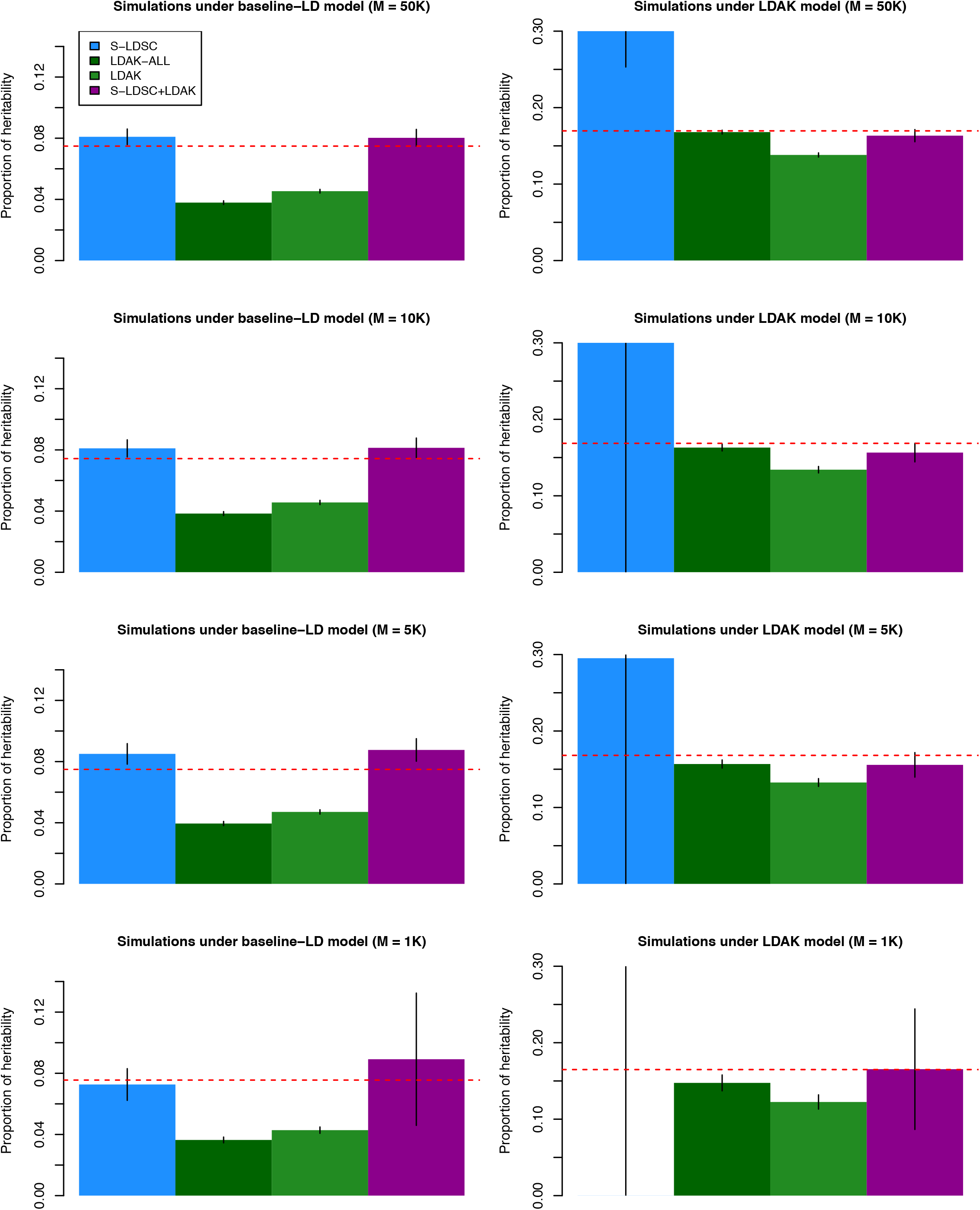
Simulations with different numbers of causal variants. We report the estimated proportion of heritability explained by coding variants in simulations in which we simulated effect sizes based on the baseline-LD model (left) or on coding enrichment under the LDAK-annotation model (right), using 50K causal variants (first row), 10K causal variants (second row), 5K causal variants (third row), and 1K causal variants (last row). The S-LDSC method uses S-LDSC with the baseline-LD model. The LDAK-allSNPs method uses the LDAK method with all the SNPs with MAF ≥ 1%. The LDAK method uses all the SNPs with MAF ≥ 1% and an INFO score ≥ 0.99 (as recommended). The S-LDSC+LDAK method uses S-LDSC with the baseline-LD+LDAK-annotation model. We report the proportion of heritability explained rather than enrichment due to the different LDAK and S-LDSC enrichment estimands. Dashed red lines indicate the true simulated value. Results are averaged across 500 simulations. Error bars represent 95% confidence intervals.

**Figure S7:**
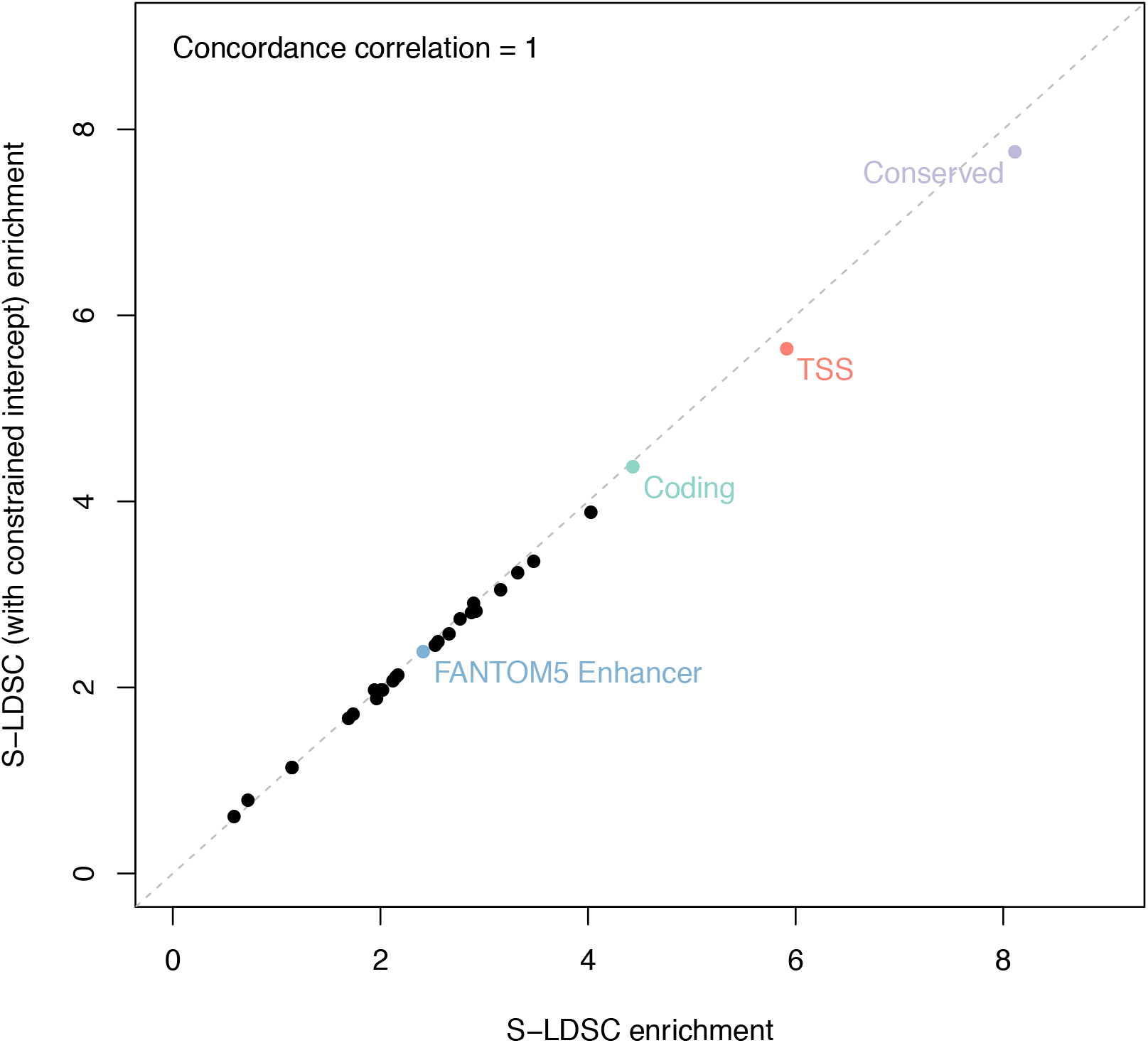
Comparison of enrichment estimates for S-LDSC vs. S-LDSC with constrained intercept. We report enrichment estimates of S-LDSC vs. S-LDSC with constrained intercept for 28 functional annotations, meta-analyzed across 16 independent UK Biobank traits. We report the concordance correlation coefficient *ρ_c_*. Grey lines represent *y* = *x*. We observed that S-LDSC results were similar to results of S-LDSC with constrained intercept (*ρ_c_* = 1.00), unlike in simulations under the LDAK model (Figure S5).

**Figure S8:**
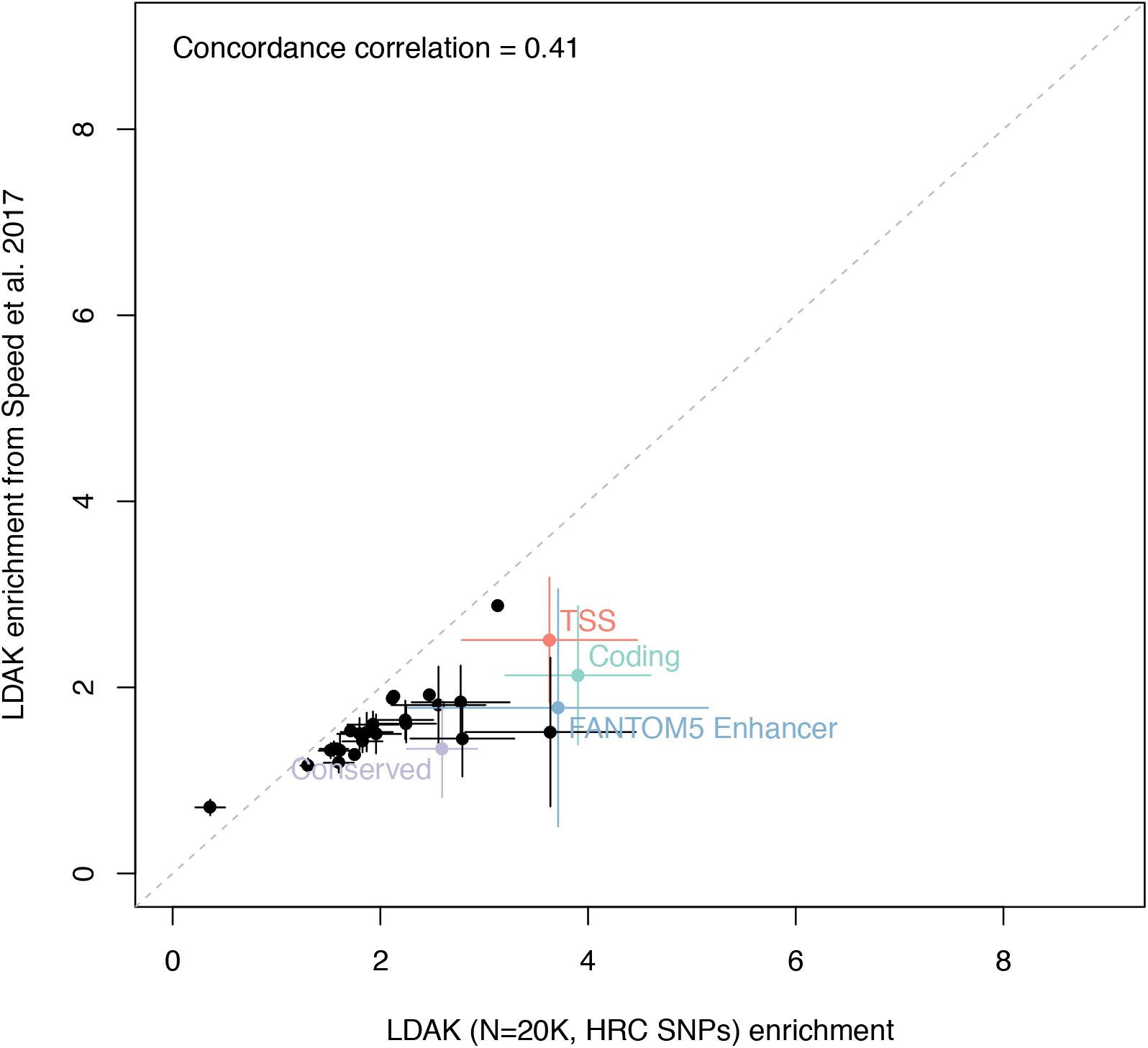
Comparison of LDAK enrichment estimates in UK Biobank data and LDAK enrichment estimates from Speed et al. 2017. We report our LDAK enrichment estimates, meta-analyzed across 16 independent UK Biobank traits, vs. LDAK enrichment estimates reported by Speed et al., for 28 functional annotations. We report the concordance correlation coefficient *ρ_c_*. Grey lines represent *y* = *x*. Error bars represent 95% confidence intervals for annotations for which the estimated enrichment is significantly different (*P* < 0.05) between the two methods. We observed that our LDAK maximum enrichment (coding regions, 3.90x, s.e. = 0.36x), was higher than the maximum significant LDAK enrichment reported in ref. ^3^ (i.e. 2.51x). We hypothesize that the difference is due to the set of traits investigated.

**Figure S9:**
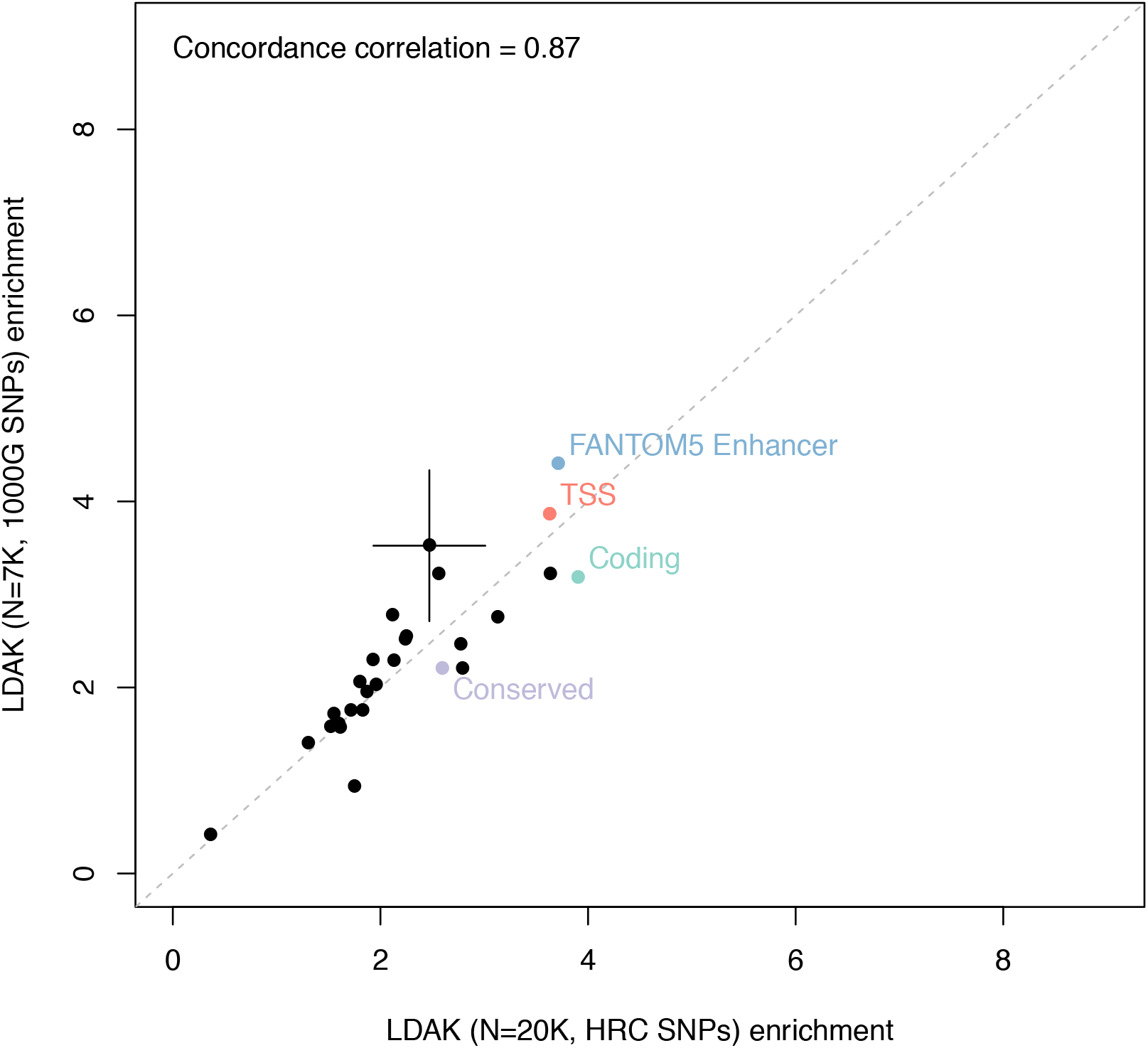
Comparison of LDAK enrichment estimates with different sample size and SNPs set. We report the enrichment of LDAK with our main setting (*N* = 20K and *M* = 4.6M HRC SNPs) vs. LDAK with similar sample size and SNPs set than Speed et al. 2017 (*N* = 7K and *M* = 2.8M 1000G SNPs) for 28 functional annotations, meta-analyzed across 16 independent UK Biobank traits. We report the concordance correlation coefficient *ρ_c_*. Grey lines represent *y* = *x*. Error bars represent 95% confidence intervals for annotations for which the estimated enrichment is significantly different (*P* < 0.05) between the two methods. We observed similar LDAK enrichment estimates for the two data sets.

**Figure S10:**
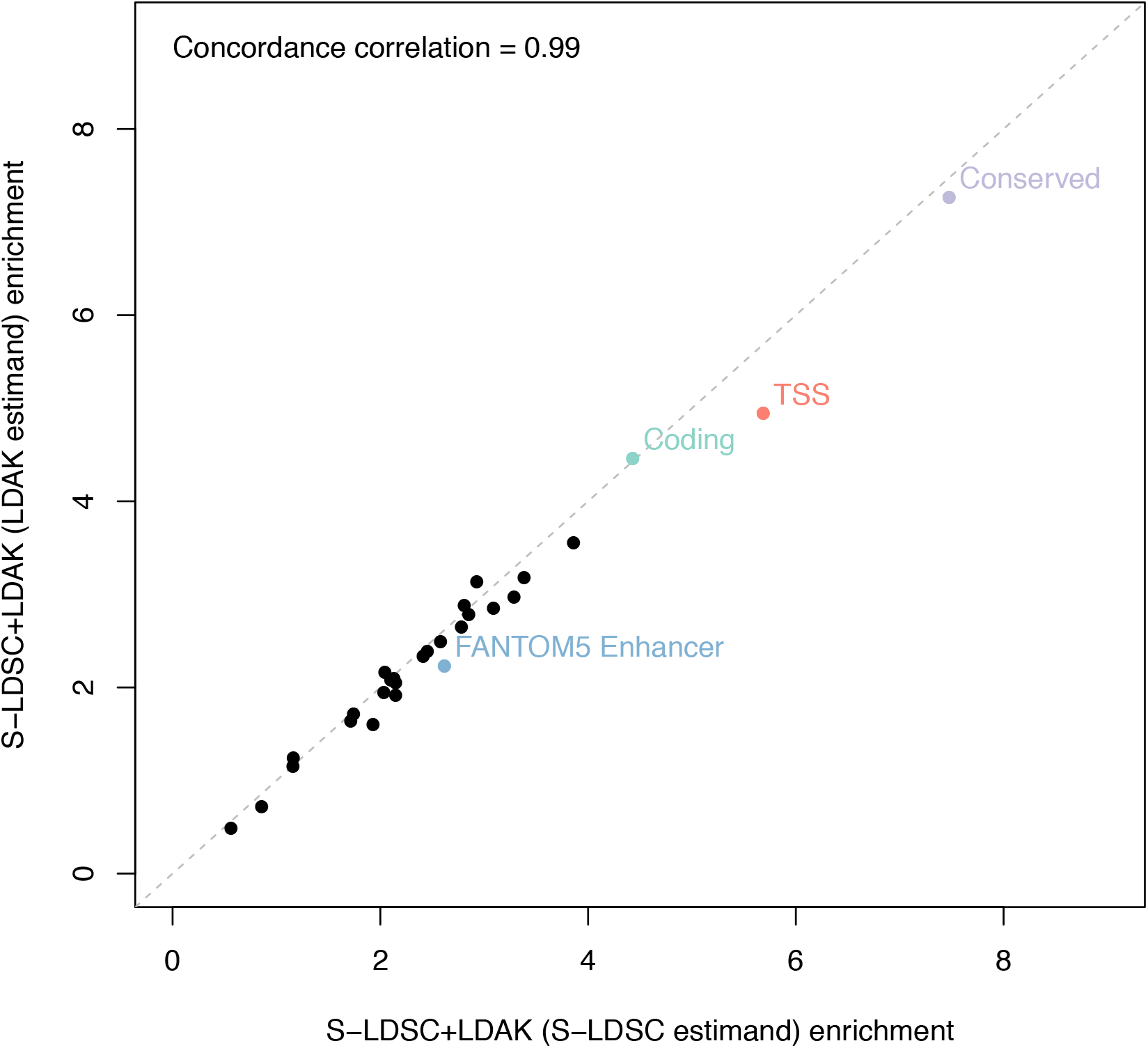
Comparison of S-LDSC+LDAK enrichment estimates using different estimands. We report the enrichment estimates of S-LDSC+LDAK (S-LDSC estimand) vs. S-LDSC+LDAK (LDAK estimand) for 28 functional annotations, meta-analyzed across 16 independent UK Biobank traits. We report the concordance correlation coefficient *ρ_c_*. Grey lines represent *y* = *x*. We observed similar results with the two different estimands.

**Figure S11:**
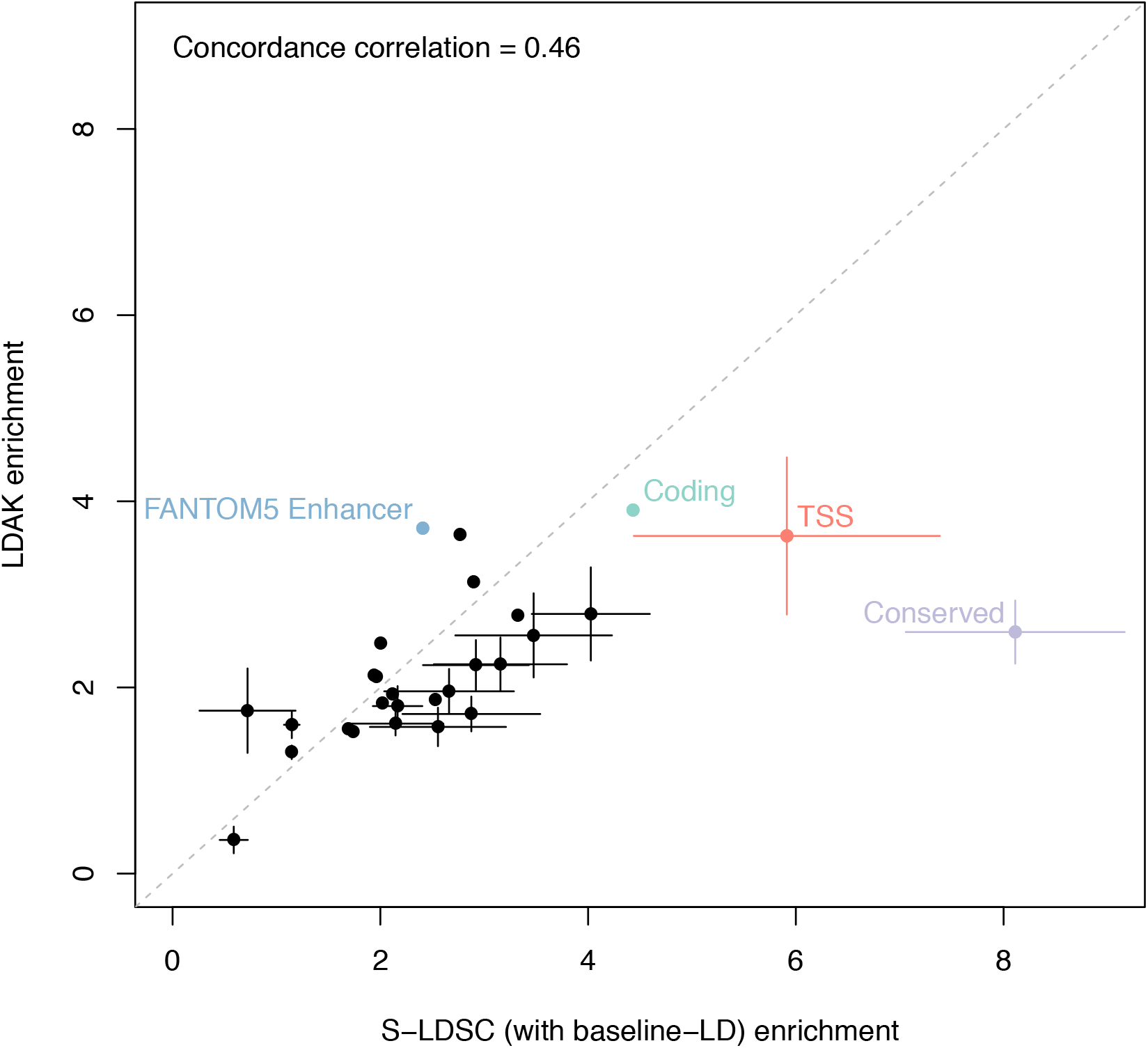
Comparison of S-LDSC and LDAK enrichment estimates. We report the enrichment estimates of S-LDSC (S-LDSC estimand) vs. LDAK (LDAK estimand) for 28 functional annotations, meta-analyzed across 16 independent UK Biobank traits. We report the concordance correlation coefficient *ρ_c_*. Grey lines represent *y* = *x*. Error bars represent 95% confidence intervals for annotations for which the estimated enrichment is significantly different (*P* < 0.05) between the two methods. We observed that LDAK enrichment estimates were higher than in ref. ^3^, with estimates closer to S-LDSC except for conserved regions.

**Figure S12:**
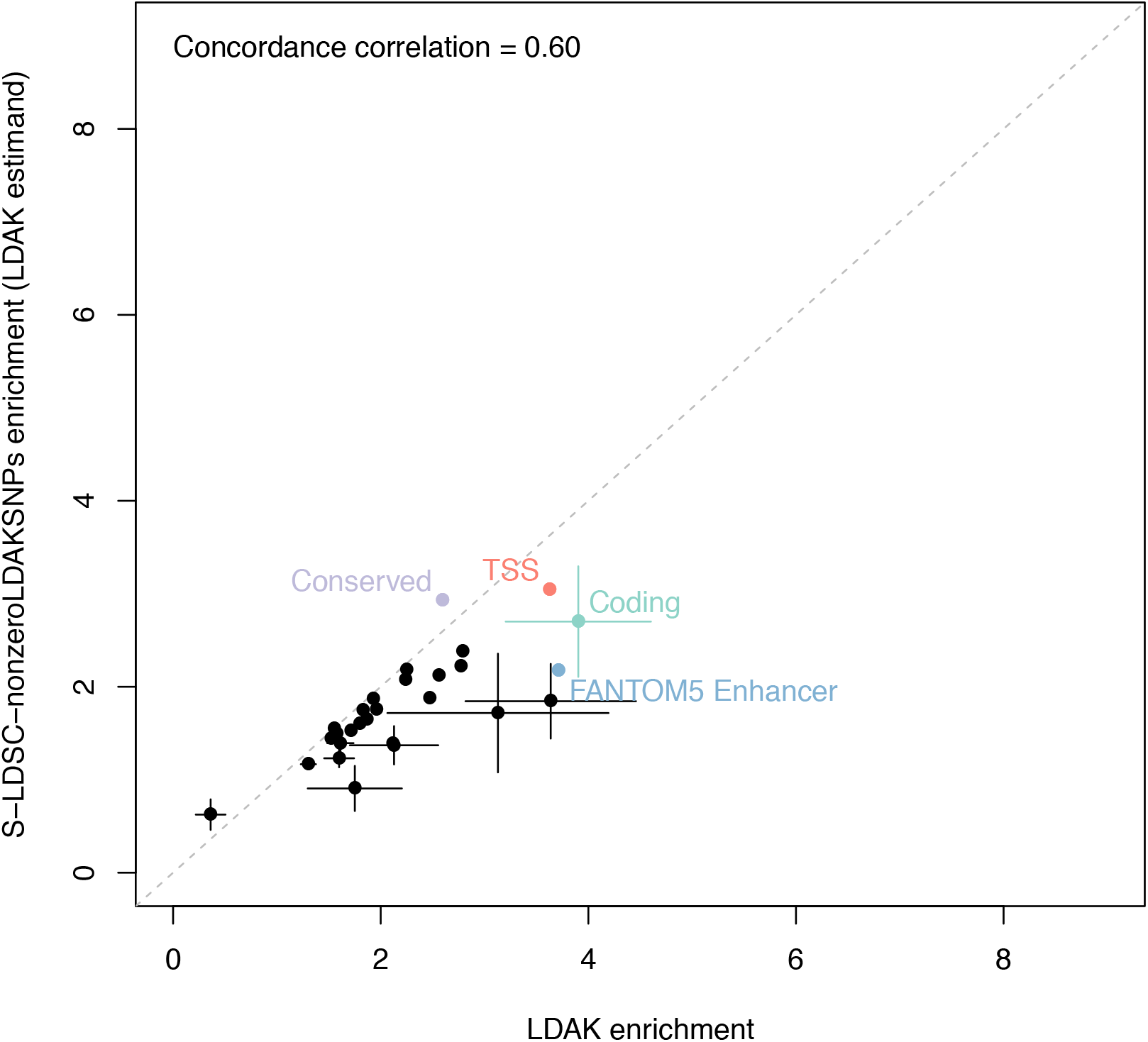
Comparison of enrichment estimates for LDAK and S-LDSC-nonzeroLDAKSNPs. We report the enrichment estimates of LDAK vs. S-LDSC-nonzeroLDAKSNPs for 28 functional annotations, meta-analyzed across 16 independent UK Biobank traits. S-LDSC-nonzeroLDAKSNPs is a method using S-LDSC with the baseline-LD model (with the LDAK estimand) that restricts the LD reference panel to the 708K SNPs with an LDAK weight different from 0 in the LDAK analyses (see Table S5). We report the concordance correlation coefficient *ρ_c_*. Grey line represents *y* = *x*. Error bars represent 95% confidence intervals for annotations for which the estimated enrichment is significantly different (*P* < 0.05) between the two methods. We observed that S-LDSC-nonzeroLDAKSNPs enrichment estimates were generally similar to or lower than LDAK estimates, with a similar enrichment estimate for the conserved annotation (2.93x, s.e. = 0.12x and 2.60x, s.e. = 0.17x, respectively).

**Figure S13:**
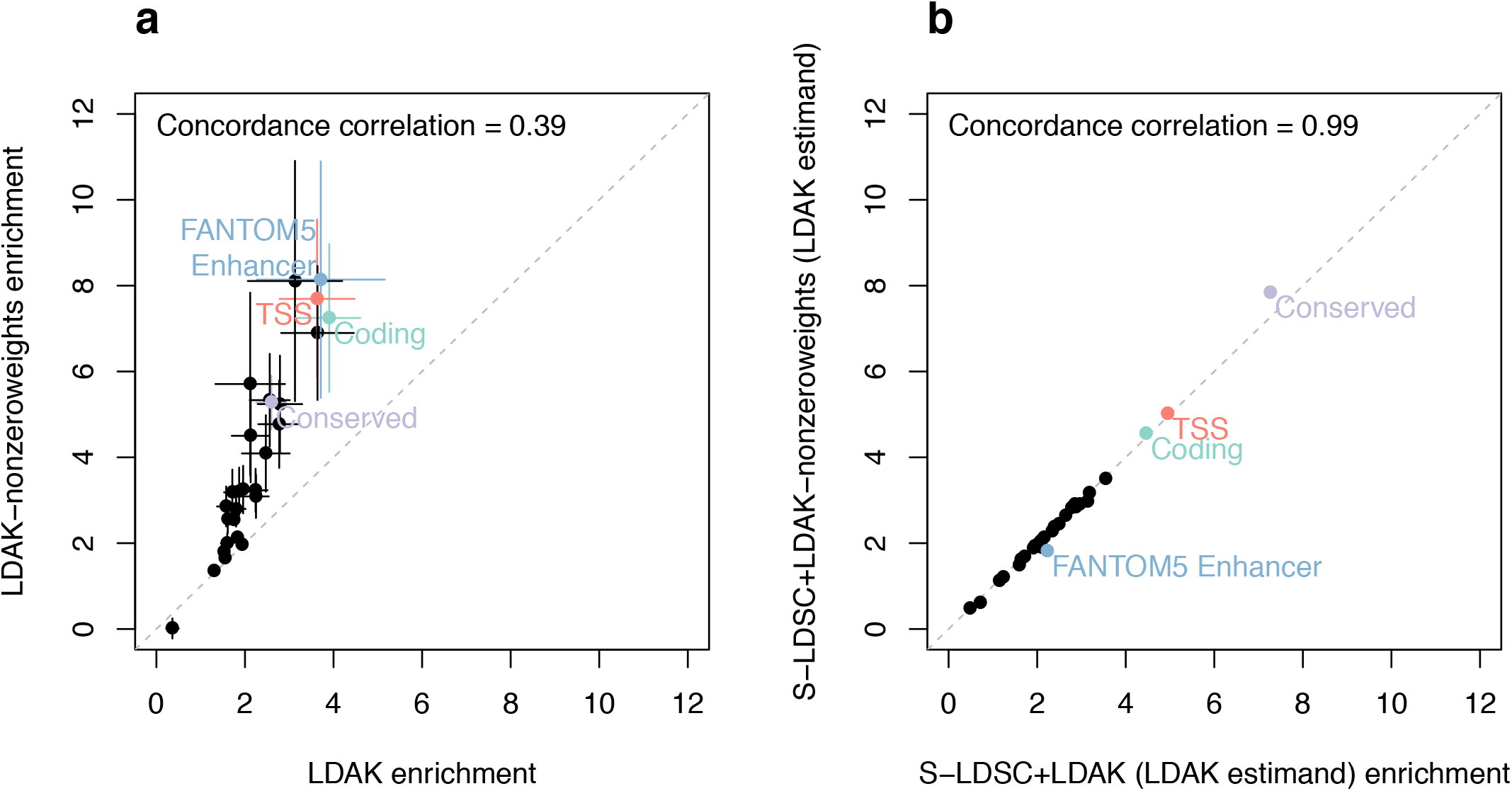
Using non-zero LDAK weights increases LDAK enrichment estimates but does not change S-LDSC+LDAK (LDAK estimand) estimates. We report the enrichment estimates of LDAK vs. LDAK-nonzeroweights (**a**) and S-LDSC+LDAK (LDAK estimand) vs. S-LDSC+LDAK-nonzeroweights (LDAK estimand) (**b**) for 28 functional annotations, meta-analyzed across 16 independent UK Biobank traits. LDAK-nonzeroweights is a method that runs LDAK software in a non-default mode to produce non-zero weights for every SNP (see Table S5). S-LDSC+LDAK-nonzeroweights (LDAK estimand) is a method analogous to S-LDSC+LDAK (LDAK estimand) that uses the non-zero weights from LDAK-nonzeroweights (see Table S5). We report the concordance correlation coefficient *ρ_c_*. Grey lines represent *y* = *x*. Error bars represent 95% confidence intervals for annotations for which the estimated enrichment is significantly different (*P* < 0.05) between the two methods.

**Figure S14:**
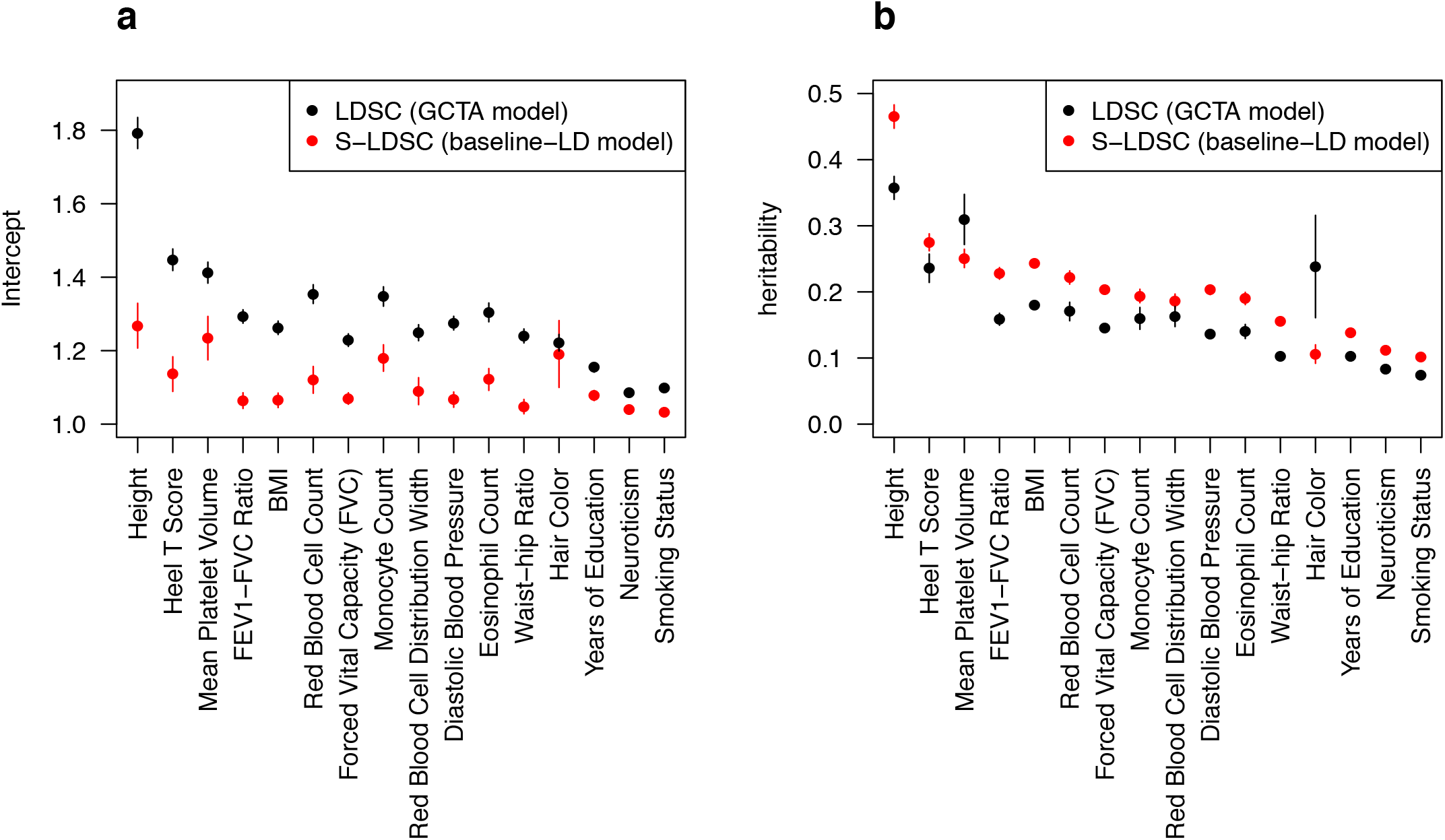
S-LDSC with the baseline-LD model produces lower regression intercepts and higher heritability estimates than LDSC. We report the regression intercepts (**a**) and estimates of the heritability causally explained by all common SNPs (**b**) for 16 independent UK Biobank traits when using S-LDSC with the baseline-LD model and LDSC, respectively. These results for regression intercept extend naturally to the attenuation ratio, a measure of calibration^19^ (defined as (regression intercept −1)/(mean *χ*^2^-1)).

**Figure S15:**
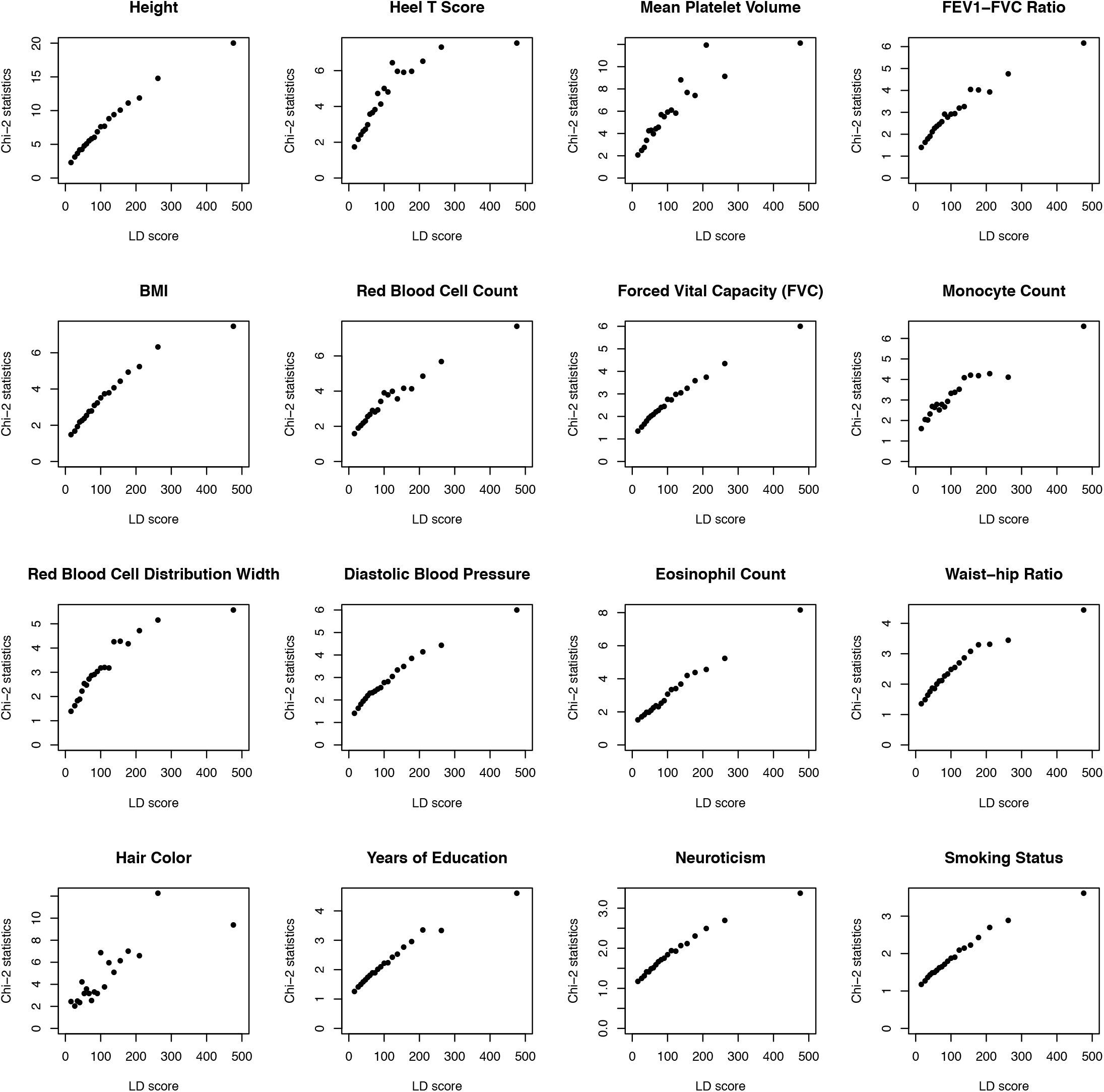
Relationship between LD scores and *χ*^2^ statistics in 16 independent UK Biobank traits. For each of 20 LD score bins, we report the mean LD scores (computed on UK10K) and the mean *χ*^2^ statistics for 16 independent UK Biobank traits (average *N* = 434K). Plots are restricted to HapMap 3 SNPs used by S-LDSC to perform the regression. The analogous plots for simulations under the LDAK and baseline-LD models are provided in Figure S1.

**Figure S16:**
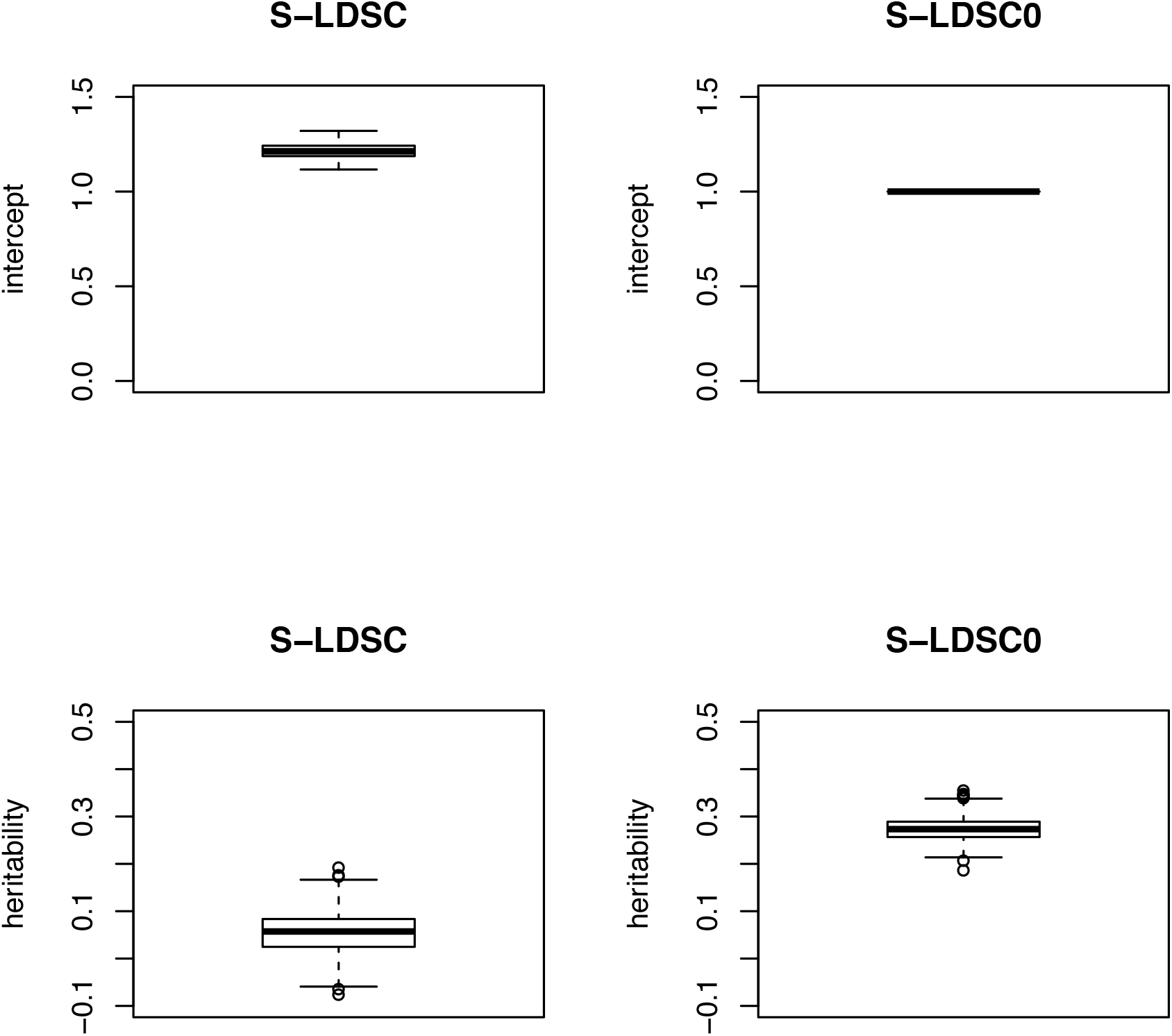
Regression intercepts and heritability estimates of S-LDSC and S-LDSC with constrained intercept (S-LDSC0) in simulations under the LDAK model. We report regression intercepts (first row) and estimates of the heritability causally explained by all common SNPs (second row) of S-LDSC (first column) and S-LDSC with constrained intercept (S-LDSC0) (second column) in 500 simulations under the LDAK model. We observed that S-LDSC (first column) produces high regression intercepts (mean value = 1.21) and small heritability estimates (mean value = 0.05). For 11% of the simulations we observed negative estimates, which creates unstable estimates of functional enrichment (see Figure S5). When constraining the intercept to 1 (S-LDSC0; right column), we observed stable heritability estimates (mean value = 0.27; no negative estimates).

**Figure S17:**
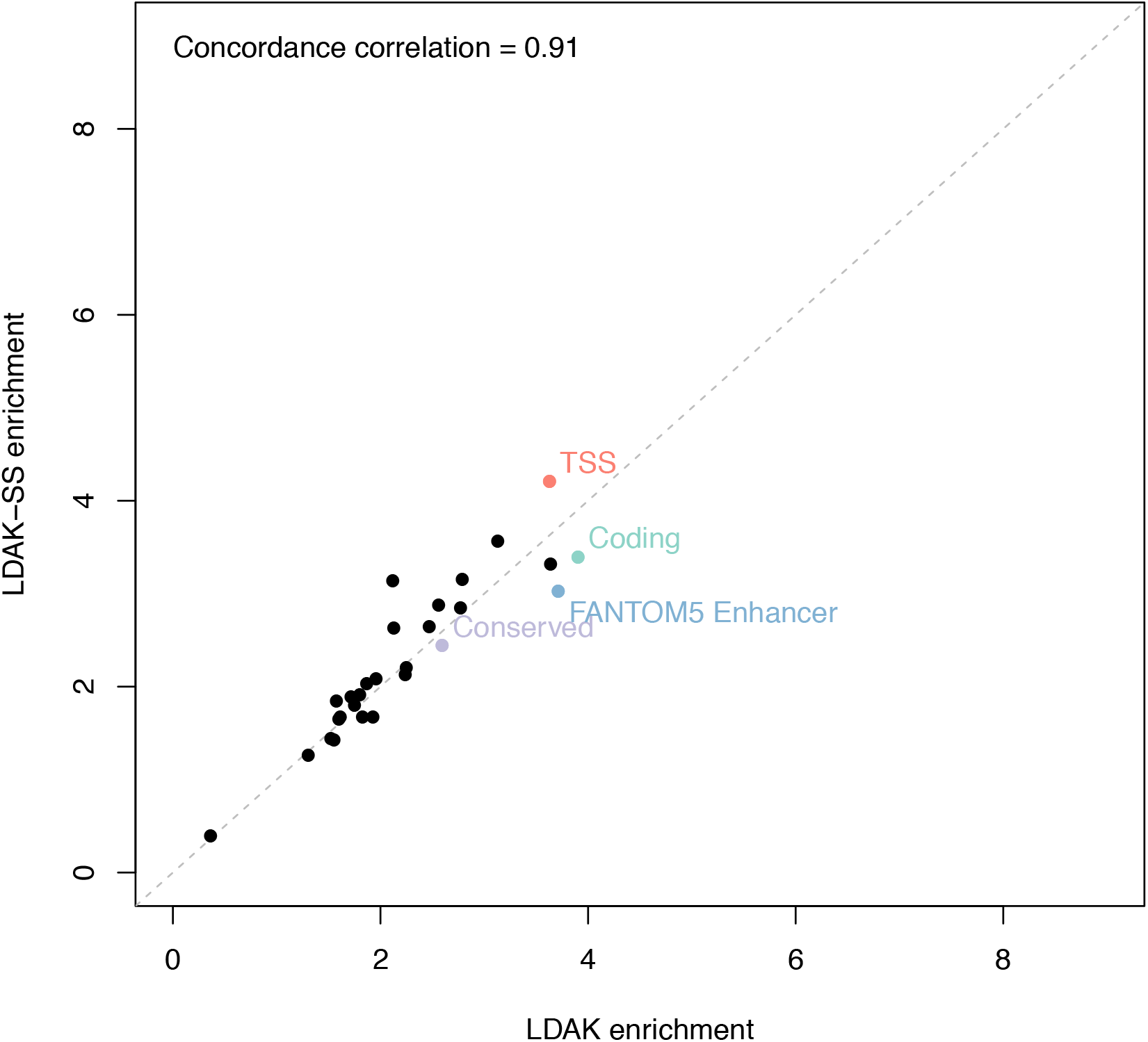
Comparison of LDAK and LDAK-SS enrichment estimates. We report the enrichment estimates of LDAK vs. LDAK-SS for 28 functional annotations, meta-analyzed across 16 independent UK Biobank traits. We report the concordance correlation coefficient *ρ_c_*. Grey line represents *y* = *x*. Error bars represent 95% confidence intervals for annotations for which the estimated enrichment is significantly different (*P* < 0.05) between the two methods. We observed similar estimates (*ρ_c_* = 0.91, no annotation significantly different), indicating that LDAK low enrichment estimates are not due to the different inference method or different sample size.

**Figure S18:**
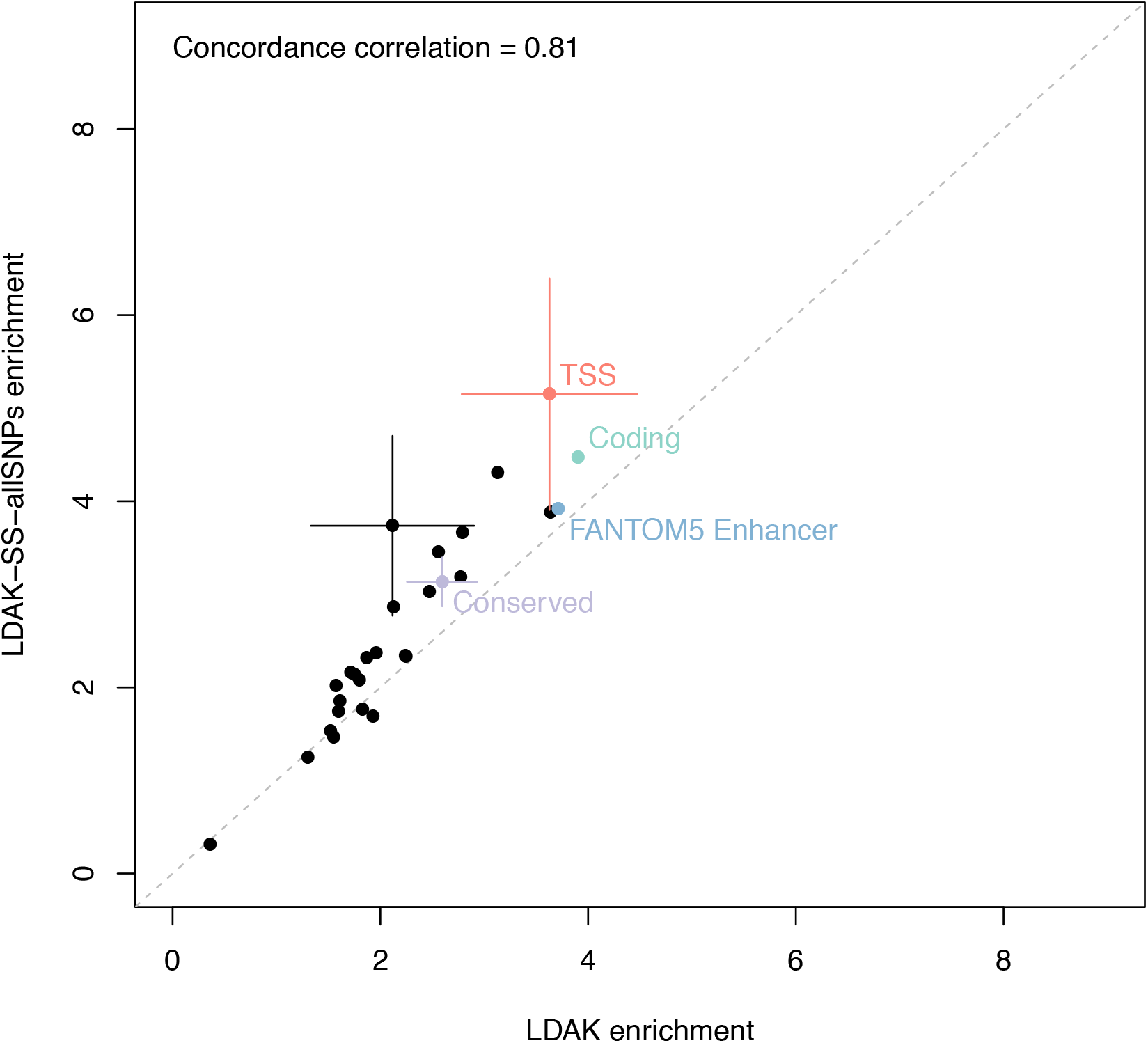
Comparison of LDAK and LDAK-SS-allSNPs enrichment estimates. We report the enrichment estimates of LDAK vs. LDAK-SS-allSNPs for 28 functional annotations, meta-analyzed across 16 independent UK Biobank traits. We report the concordance correlation coefficient *ρ_c_*. Grey line represents *y* = *x*. Error bars represent 95% confidence intervals for annotations for which the estimated enrichment is significantly different (*P* < 0.05) between the two methods. We observed low conserved enrichment estimates when using LDAK-SS-allSNPs (3.13x, s.e. = 0.13x), indicating that low conserved enrichment estimates for LDAK (and LDAK-SS) are not due to the use of well-imputed SNPs.

**Figure S19:**
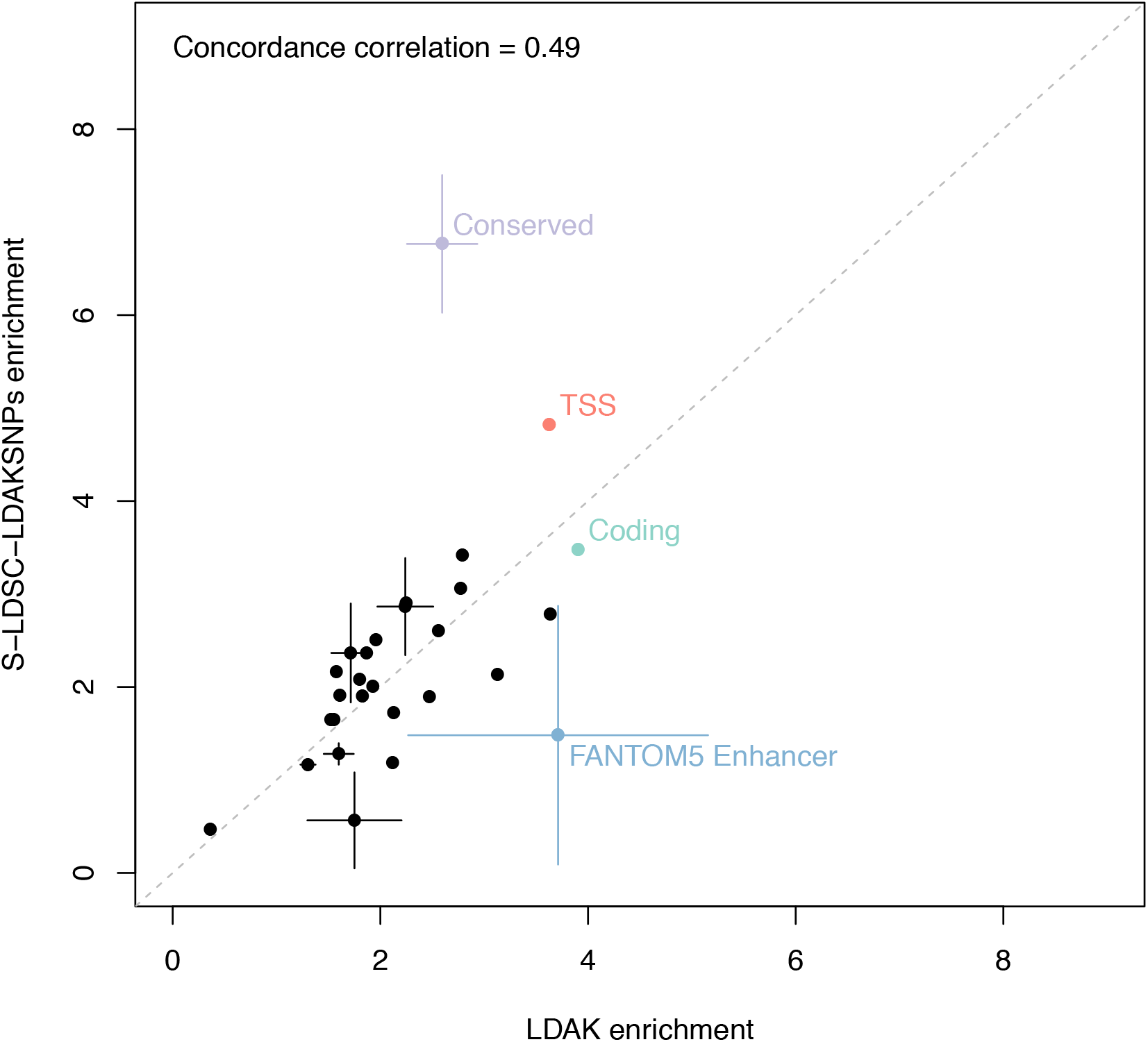
Comparison of LDAK and S-LDSC-LDAKSNPs enrichment estimates. We report the enrichment estimates of LDAK vs. S-LDSC-LDAKSNPs for 28 functional annotations, meta-analyzed across 16 independent UK Biobank traits. We report the concordance correlation coefficient *ρ_c_*. Grey line represents *y* = *x*. Error bars represent 95% confidence intervals for annotations for which the estimated enrichment is significantly different (*P* < 0.05) between the two methods. We observed high conserved enrichment estimates for S-LDSC-LDAKSNPs (6.77x, s.e. = 0.38x), indicating that high S-LDSC enrichment estimates (compared to LDAK) are not due to the use of well-imputed SNPs.

